# SUMO orchestrates multiple alternative DNA-protein crosslink repair pathways

**DOI:** 10.1101/2020.06.08.140079

**Authors:** Nataliia Serbyn, Ivona Bagdiul, Agnès H. Michel, Raymond T. Suhandynata, Huilin Zhou, Benoît Kornmann, Françoise Stutz

**Affiliations:** Department of Cell Biology, University of Geneva, 1211 Geneva 4, Switzerland; Institute of Biochemistry, ETH Zürich, 8093 Zurich, Switzerland; Department of Biochemistry, University of Oxford, Oxford OX1 3QU, United Kingdom; Department of Cellular and Molecular Medicine, Moores Cancer Center, University of California School of Medicine, San Diego, La Jolla, United States

**Keywords:** SUMO, DNA-protein crosslink, DPC repair, Top1, Top1cc, Siz2, Ulp1, Wss1, Tdp1, STUbL, Flp-nick

## Abstract

Several endogenous metabolites, environmental agents, and therapeutic drugs promote formation of covalent DNA-protein crosslinks (DPCs). Persistent DPCs pose a serious threat to genome integrity and are eliminated by multiple repair pathways. Aberrant Top1 crosslinks to DNA, or Top1ccs, are processed by Tdp1 and Wss1 functioning in parallel pathways in *Saccharomyces cerevisiae.* It remains obscure how cells choose between these diverse mechanisms of DPC repair. Here we show that several SUMO biogenesis factors - Ulp1, Siz2, Slx5, Slx8 - control repair of Top1cc or an analogous DPC lesion. Genetic analysis reveals that SUMO promotes Top1cc processing in the absence of Tdp1 but has an inhibitory role if cells additionally lack Wss1. In the *tdp1Δ wss1Δ* mutant, the E3 SUMO ligase Siz2 stimulates sumoylation in the vicinity of the DPC, but not SUMO conjugation to Top1. This Siz2-dependent sumoylation delays DPC repair when cells progress through S and G2 phases. Our findings suggest that SUMO tunes available repair pathways to facilitate faithful DPC repair.

## INTRODUCTION

The interaction between two major biopolymers – DNA and protein – is indispensable for the storage and flow of genetic information. DNA-protein complexes are typically dissociated in the order of seconds (Phair et al., 2004), while formation of an aberrant covalent bond abrogates this process. Persistent DNA-protein crosslinks (DPCs) are highly toxic if not resolved in a timely manner, as they cause genome instability and eventually promote cell death (Ide et al., 2018; Klages-Mundt and Li, 2017; Stingele et al., 2017).

Several classes of enzymes form a covalent bond with DNA as part of their catalytic mechanism. One example is DNA topoisomerases that perform DNA cleavage to rotate DNA and relax the torsional stress arising from replication, transcription and chromatin remodeling (Pommier et al., 2016). Yeast topoisomerase 1 (Top1) nicks a single DNA strand to form a transient covalent reaction intermediate with the DNA 3’ end. If the subsequent DNA sealing step cannot be completed, Top1 remains trapped on DNA in a covalent cleavage complex (Top1cc). Top1 trapping can occur spontaneously, especially in the presence of a proximal DNA lesion. Top1cc stabilization can also be induced by natural small molecules, e.g. camptothecin (CPT), or pharmaceutically developed topoisomerase inhibitors (Pommier, 2006). CPT and its derivatives enter the catalytic pocket of Top1, thus preventing the DNA re-ligation step and stabilizing a DPC on the nicked 3’ end of DNA (Staker et al., 2002). Several Top1 inhibitors are used in chemotherapeutics, as the excessive Top1ccs collide with replication forks, generate DNA damage, and cause cytotoxicity for actively dividing cancer cells (Hengel et al., 2017; Pommier, 2006).

Numerous enzymes and repair pathways were previously implicated in Top1cc processing. Eukaryotic cells carry a specialized enzyme that directly hydrolyzes the Top1-DNA covalent bond - the phosphodiesterase TDP1 (Kawale and Povirk, 2018). Alternatively, Top1ccs can be excised from DNA by several nuclease complexes such as yeast Rad1-Rad10 (human XPF-ERCC1), Slx4-Slx1 (hSLX4-SLX1), Mus81-Mms4 (hMUS81-EME1), Mre11-Rad50-Xrs2 (hMRE11-RAD50-NBS1), and Rad27 (hFEN1) (reviewed in (Sun et al., 2020)). Importantly, large proteins such as full-length Top1 are poor *in vitro* TDP1 substrates and require at least a partial proteolysis (Debethune et al., 2002). Degradation of the DPC protein moiety can be achieved either by the canonical 26S proteasome machinery (Desai et al., 2001; Larsen et al., 2019; Lin et al., 2008; Zhang et al., 2004), or by the DPC-specific metalloprotease Wss1 in yeast or SPRTN in higher eukaryotes (Lopez-Mosqueda et al., 2016; Maskey et al., 2017; Stingele et al., 2016; Stingele et al., 2014; Vaz et al., 2016). Several other proteases were proposed to function in DPC proteolysis – GCNA, a mammalian SPRTN homologue (Bhargava et al., 2020; Borgermann et al., 2019; Dokshin et al., 2020), the serine protease FAM111A (Kojima et al., 2020), as well as the yeast aspartic protease Ddi1 (Serbyn et al., 2020). The Cdc48 molecular segregase (p97 in mammals) also assists the Top1cc extraction process (Fielden et al., 2020; Nie et al., 2012; Stingele et al., 2014). Following partial proteolysis, Top1 peptides that remain attached to the single stranded (ss) DNA break can be further removed by the canonical DNA repair pathways such as nucleotide or base excision repair (NER and BER) involving the aforementioned nucleases and TDP1, as well as PARP1, PNKP, polymerase *β*, and DNA ligase (Mei et al., 2020; Pommier et al., 2006). If a ssDNA break persists until S-phase, collision of the replication fork with the Top1cc may generate a one-ended double-stranded break (DSB) that can be repaired by homologous recombination (HR) (Pommier et al., 2003).

Mechanisms orchestrating the DPC repair pathway choice remain, however, poorly understood. Emerging evidence indicates that posttranslational modifications (PTMs) such as phosphorylation, ubiquitination and sumoylation can play a pivotal role in both DPC recognition and processing (Borgermann et al., 2019; Das et al., 2009; Desai et al., 2000; Duxin et al., 2014; Huang et al., 2010; Larsen et al., 2019). Among them, conjugation of the Small Ubiquitin-like MOdifier (SUMO) to Top1 was previously reported both in yeast and mammalian cells (Chen et al., 2007; Desai et al., 2000; Kanagasabai et al., 2009; Mao et al., 2000b). Whereas its function in yeast remains enigmatic, CPT-induced Top1 sumoylation in higher eukaryotes promotes Top1 exclusion from the highly transcribed rDNA locus (Mao et al., 2000b; Mo et al., 2002). SUMO can also be conjugated to several repair factors involved in Top1cc processing including the nucleases SLX4, FEN1 and yeast Rad1 (Ouyang et al., 2015; Sarangi et al., 2014; Xu et al., 2018), the recombinase Rad52 (Sacher et al., 2006), the non-homologous end-joining (NHEJ) protein Yku70 (Hang et al., 2014), and PCNA (Pfander et al., 2005). Moreover, efficient DSB repair is tuned by “on-site” sumoylation, which involves multiple HR repair factors (Cremona et al., 2012; Psakhye and Jentsch, 2012).

SUMO conjugation is generally thought to promote the efficient processing of Top1ccs and other DPC types (Borgermann et al., 2019; Chen et al., 2007; Jacquiau et al., 2005; Mao et al., 2000b; Nie et al., 2012; Schellenberg et al., 2017). However, several studies suggest the opposite: SUMO conjugation appears to have a deleterious effect on Top1cc repair (Horie et al., 2002; Sharma et al., 2017). In this work, we show that SUMO conjugation can indeed have a dual role in the Top1cc repair in *Saccharomyces cerevisiae*. We found that localization of the Ulp1 SUMO protease must be restricted to the nuclear pore complex in order to maintain normal protein levels of the two E3 SUMO ligases – Siz1 and Siz2. Among these, the Siz2 E3 SUMO ligase promotes sumoylation at the DPC site but does not target Top1 itself. Sumoylation by Siz2 aids Top1cc repair in the absence of Tdp1 but becomes deleterious if the Wss1 protease is additionally unavailable. The excessive on-site sumoylation by Siz2 in *tdp1Δ wss1Δ* inhibits DPC repair when cells progress through the S and G2 cell cycle phases. In summary, we propose that cell cycle dependent sumoylation controls the processing of DNA-protein crosslinks such as Top1ccs.

## RESULTS

### A transposon screen identifies multiple components of the SUMO biogenesis pathways as suppressors of *tdp1 wss1*

Top1-DNA crosslinks accumulate in the *tdp1Δ wss1Δ* yeast cells lacking the two key Top1cc repair components – the phosphodiesterase Tdp1 and the DPC protease Wss1 (Stingele et al., 2014). To gain insight into the regulatory mechanisms controlling Top1cc processing, we exploited results of a Saturated Transposon Analysis in Yeast (SATAY) performed in the *tdp1-AID wss1Δ* mutant (Serbyn et al., 2020). The *tdp1-AID* auxin-inducible degron (AID) system (Morawska and Ulrich, 2013) allows to rapidly deplete Tdp1 and permits studying the nearly unviable *tdp1Δ wss1Δ* double deletion mutant. To identify mutations that suppress synthetic sickness of the double mutant, we compared transposition events in *tdp1-AID wss1Δ* + auxin and several WT-like libraries (Figure 1A) using the “read_per_gene” value best suited to identify suppressor mutations (Michel et al., 2017). This analysis identified *TOP1* among the strongest *tdp1-AID wss1Δ* suppressors (Figure 1A), supporting previous reports that *tdp1Δ wss1Δ* synthetic growth defect is mainly due to unrepaired Top1css (Balakirev et al., 2015; Stingele et al., 2014). Among other *tdp1-AID wss1Δ* suppressors were several components of the nuclear pore complex (NPC) (Figure 1A, highlighted in green). We confirmed that deletion of *NUP60* indeed rescues the growth phenotype of *tdp1Δ wss1Δ* (Figure 1B). Remarkably, all identified nucleoporins are present at the nuclear side of the NPC (Figure S1A). In the same screen, SUMO biogenesis components were also enriched among *tdp1-AID wss1Δ* + auxin suppressors (Figure 1A, highlighted in red). In yeast, nuclear pore basket proteins Mlp1, Mlp2, as well as Nup60 and the Nup84 complex, anchor the Ulp1 SUMO protease to the NPC (Palancade et al., 2007; Zhao et al., 2004) (Figure S1A); we therefore hypothesized that lack of Ulp1 tethering to the NPC promotes *tdp1Δ wss1Δ* survival. To test this possibility, we took advantage of the fact that the N-terminal domain deletion mutant *ulp1-ΔN* (Figure 1C) de-localizes the SUMO protease to the nucleoplasm (Texari et al., 2013). Consistent with our hypothesis, Ulp1 de-localization was sufficient to suppress *tdp1Δ wss1Δ* (Figure 1D). The suppression was dominant (Figure S1B) and required the catalytically active protease domain of Ulp1 (Figures 1C and 1E), indicating that the suppression is caused by Ulp1 desumoylation activity in the nucleoplasm and not by its absence from NPCs.

**Figure 1.**
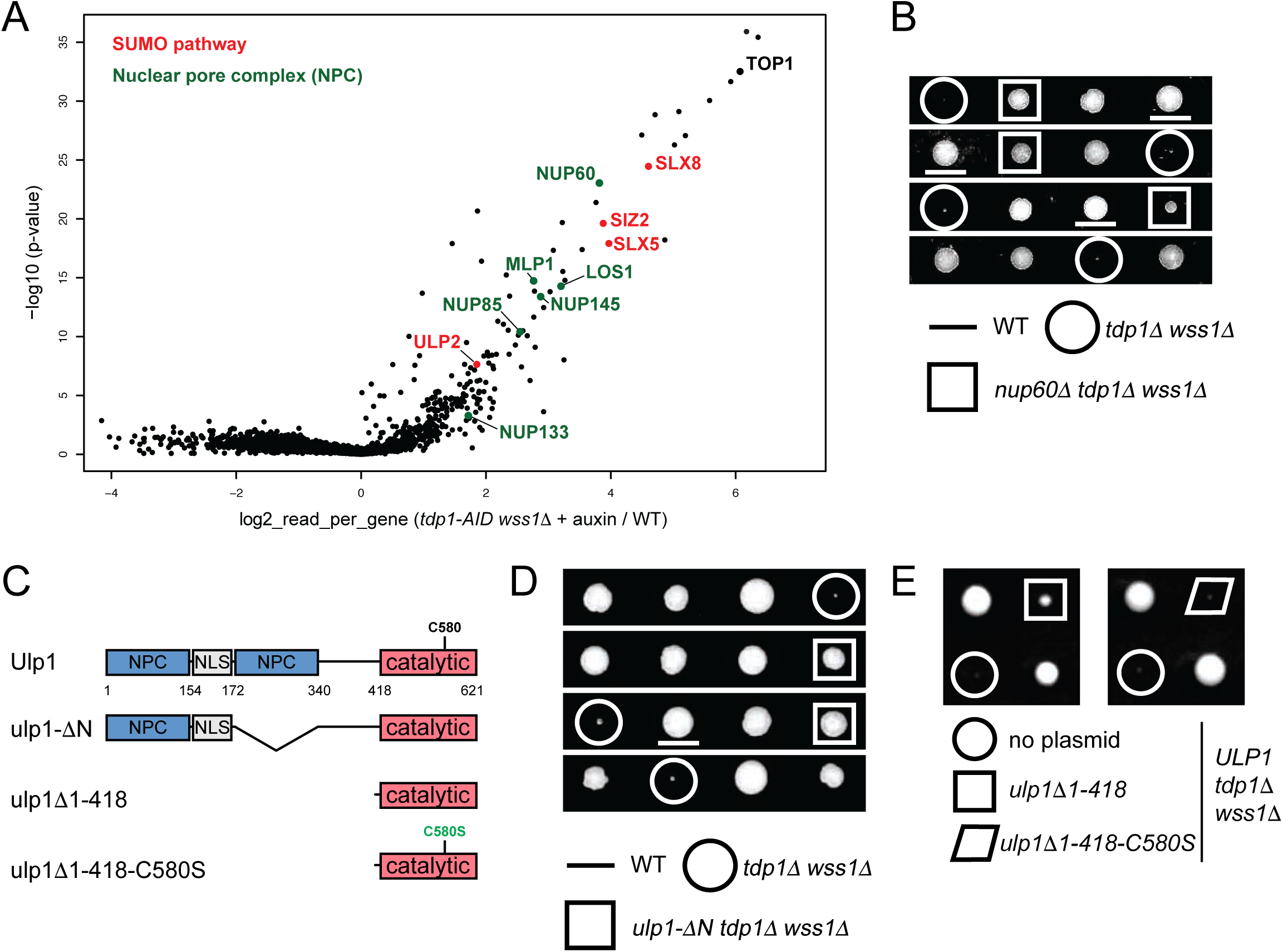
Mutants of nucleoporins and SUMO biogenesis factors act as suppressors of the *tdp1 wss1* double mutant phenotypes (A) Suppressors identified in the *tdp1-AID wss1Δ* genetic screen described in (Serbyn et al., 2020). *tdp1-AID* is the shortening of [*TDP1-AID*-6HA; pADH-TIR1*]. The volcano plot compares sequencing reads in *tdp1-AID wss1Δ*+auxin and a pool of unrelated SATAY libraries. Fold-changes of reads per gene (log2, x-axis) and corresponding p-values (- log10, y-axis) are plotted. SUMO-related genes are highlighted in red. Genes that encode nucleoporins and nuclear pore complex-associated proteins are shown in green. (B) Loss of the Nup60 nucleoporin suppresses the growth defect of *tdp1*Δ *wss1*Δ. [*TDP1/tdp1Δ; WSS1/wss1Δ; NUP60/nup60Δ*] diploid was used for the tetrad analysis. These and subsequent selected tetrads presented in the same panel were grown on the same plate. (C) A schematic illustration of the Ulp1 SUMO protease and its mutant variants. NPC, region important for anchoring to the nuclear pore complex; NLS, nuclear localization signal; catalytic, the domain responsible for SUMO maturation and cleavage; C580S, the catalytic active site mutation; the scheme is based on (Li and Hochstrasser, 2003). The *ulp1-*Δ*N* mutant (Texari et al., 2013) additionally contains in-frame GFP that replaces amino acids 172-340 (not shown). (D) The *ulp1-*Δ*N* mutation in Ulp1 N-terminal domain strongly suppresses the *tdp1*Δ *wss1*Δ growth defect. Tetrad analysis of the [*TDP1/tdp1Δ; WSS1/wss1Δ; ULP1/ ulp1-*Δ*N*] diploid. (E) The suppression effect of Ulp1 delocalization is dominant and depends on the desumoylation activity of Ulp1. [*TDP1/tdp1*Δ*; WSS1/wss1*Δ] diploid was transformed with plasmids coding for the N-terminally truncated catalytically active (*ulp1Δ1-418*) or inactive (*ulp1Δ1-418-C580S*) Ulp1 mutants expressed from the endogenous *ULP1* promoter (Li and Hochstrasser, 2003); diploids were further analyzed by tetrad analysis. Each black square contains one tetrad.

In *S. cerevisiae,* the essential Aos1-Uba2 (E1) and Ubc9 (E2) enzymes conjugate SUMO, while the substrate specificity is defined by several mitotic E3 SUMO ligases - Siz1, Siz2, and Mms21. SUMO cleavage is catalyzed by the SUMO proteases Ulp1 and Ulp2; the former is also responsible for C-terminal SUMO cleavage required for its maturation (Johnson, 2004). SUMO-dependent ubiquitin ligases (STUbLs) bind poly-sumoylated substrates through SUMO interaction motifs (SIMs) and ubiquitinate them, thus creating mixed SUMO-Ub chains (Praefcke et al., 2012). Among suppressors of the *tdp1-AID wss1Δ* screen we identified the E3 SUMO ligase Siz2, the SUMO protease Ulp2, as well as both subunits of the Slx5-Slx8 STUbL (Figure 1A). Several of them were previously isolated through an independent genetic suppressor screen of *tdp1Δ wss1Δ*, and Slx5 was further characterized to control Top1cc processing (Sharma et al., 2017). In our hands, loss of Slx5, Slx8 (Figures S1C and S1D), or the Ulp2 SUMO protease (Figure S1E) weakly suppresses *tdp1Δ wss1Δ*. Interestingly, whereas *slx5Δ* or *slx8Δ* improved the growth of *tdp1Δ wss1Δ*, the same mutants were deleterious in combination with the single *tdp1Δ* mutation in the presence of CPT (Figure S1F and data not shown), suggesting that the Slx5-Slx8 complex may participate in the Wss1-dependent pathway of DPC processing. Together, the above genetic interactions implicate sumoylation and SUMO-dependent ubiquitination in the control of Top1cc processing.

### Ulp1 activity in the nucleoplasm promotes *tdp1*Δ *wss1*Δ survival and lowers DNA damage levels through degradation of the E3 SUMO ligase Siz2

We next analyzed genetic interactions of the E3 SUMO ligases mutants and *tdp1Δ wss1Δ*. Loss of Siz2 was not only a potent suppressor of the mutant, consistent with the genetic screen results (Figures 1A and 2A), but also provided CPT resistance to *tdp1Δ wss1Δ* (Figure S2A). In contrast, the *siz1Δ* mutation was completely inviable in combination with *tdp1Δ wss1Δ*, whereas simultaneous deletions of *SIZ1* and *SIZ2* resulted in partial suppression, and *mms21-11* had no or little effect on the double mutant (Figure 2A). Remarkably, Ulp1 de-localization to the nucleoplasm promoted severe reduction in the protein levels of Siz1 and Siz2, but not Mms21 (Figure 2B). These data suggest an explanation for the puzzling similarity in the genetic interactions of the SUMO conjugation (Siz2) and de-conjugation (Ulp1) factors despite their opposing enzymatic activities (Figures 1D and 2A): Ulp1 activity in the nucleoplasm might rescue *tdp1Δ wss1Δ* phenotypes by promoting the degradation of the Siz2 SUMO ligase. To test this idea, we set out to restore Siz2 protein levels in *ulp1-ΔN tdp1Δ wss1Δ*. Siz2 overexpression to wild-type-like levels (Figure S2B) negatively impacted growth and CPT sensitivity of the *ulp1-ΔN tdp1Δ wss1Δ* mutant (Figure 2C), supporting our hypothesis.

**Figure 2.**
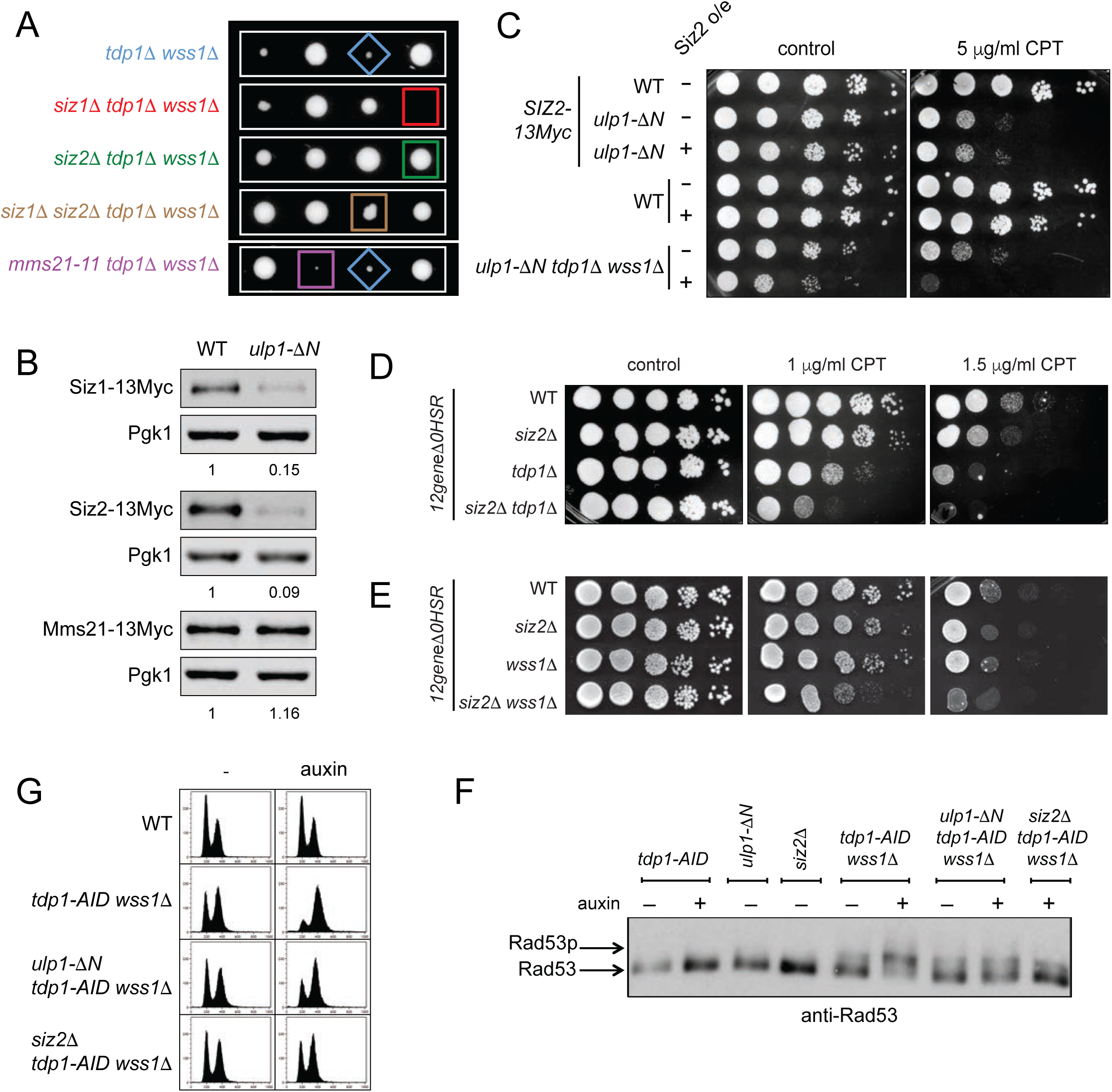
Loss of Ulp1 tethering to the nuclear periphery suppresses *tdp1*Δ *wss1*Δ via degradation of the Siz2 SUMO ligase (A) The absence of Siz2, but not Siz1 and Mms21, efficiently suppresses growth defects of *tdp1*Δ *wss1*Δ. Tetrad analysis of [*TDP1/tdp1Δ; WSS1/wss1Δ; SIZ1/siz1Δ; SIZ2/siz2Δ*] and [*TDP1/tdp1Δ; WSS1/wss1Δ; MMS21/mms21-11*] diploids. Representative spores (with the exception of the tetrad showing *mms21-11*) were grown on the same YEPD plate. (B) Siz1 and Siz2, but not Mms21 protein levels are strongly reduced in *ulp1-*Δ*N*. Cells expressing E3 SUMO ligases genomically tagged with 13Myc at their C-terminus were subjected to immunoblotting. Protein levels were normalized to Pgk1 and then to WT. (C) Decreased Siz2 protein levels in *ulp1-*Δ*N* are essential to rescue *tdp1*Δ *wss1*Δ. Strains were grown on Sc-leu to select for plasmids expressing (+) or not (-) Siz2-13Myc and imaged after 48 hours. See also Figure S2B for Siz2 protein levels. (D) Tdp1 and Siz2 provide Top1ccs resistance in a non-epistatic manner. Indicated mutations were introduced in the CPT-permeable *12geneΔ0HSR* multi-transporter mutant (Chinen et al., 2011); images were taken 72 h post-plating. (E) Wss1 and Siz2 function in the same pathway. Experiment was performed as in (D). (F) *ulp1-*Δ*N* and *siz2*Δ mutants decrease DNA damage checkpoint activation in *tdp1-AID wss1,* as measured by Rad53 phosphorylation levels by immunoblotting. Where indicated, 1 mM auxin was added for 6 h. (G) *ulp1-*Δ*N* and *siz2*Δ mutants allow progression of *tdp1-AID wss1*Δ through the cell cycle. 1 mM auxin was added for 8 h and cells were subjected to FACS analysis.

The cell wall of canonical laboratory budding yeast strains is poorly permeable to CPT. As a result, CPT fails to impair the growth of individual *tdp1Δ*, *wss1Δ*, as well as double *siz2Δ wss1Δ* and *siz2Δ tdp1Δ* mutants even at high concentrations (Figure S2C). We previously showed that the *12geneΔ0HSR* multi-transporter mutant (Chinen et al., 2011) reveals weak sensitivity of *wss1Δ* to low dosage of CPT (Serbyn et al., 2020). In the *12geneΔ0HSR* genetic background, *tdp1Δ* was also sensitive to CPT (Figure 2D). The additional *siz2Δ* mutation showed an even more pronounced growth defect on CPT (Figure 2D), suggesting that Siz2-dependent sumoylation promotes Top1cc repair independently of Tdp1. In contrast, *siz2Δ* was mostly epistatic to *wss1Δ*, suggesting that these two factors control the same aspect of Top1cc processing (Figure 2E). These genetic interactions suggest that Siz2 and Wss1 may be part of the same pathway, however the weaker CPT sensitivity of *tdp1Δ siz2Δ*, as compared to *tdp1Δ wss1Δ*, indicates that Wss1 activity could be modulated by other pathways as well.

Since the *tdp1Δ wss1Δ* strain rapidly accumulates suppressor mutations, we used the auxin-inducible Tdp1 depletion in most of subsequent experiments with the double mutant. As expected, the additional *ulp1-ΔN* or *siz2Δ* mutations suppressed *tdp1-AID wss1Δ* in the presence of auxin (Figures S2D).

The slow growth phenotype of *tdp1Δ wss1Δ* is associated with hyperactivation of the DNA damage checkpoint and the G2/M cell cycle arrest. Inactivation of the checkpoint signaling is not sufficient to rescue the growth defect (Stingele et al., 2014), suggesting that the elevated checkpoint does not promote cell death but simply reflects accumulation of unrepaired Top1ccs in *tdp1Δ wss1Δ*. We therefore reasoned that the rescue of the *tdp1Δ wss1Δ* growth defect by *ulp1-ΔN* or *siz2Δ* must be due to a decrease in Top1cc levels. Consistent with this hypothesis, auxin treatment activated the checkpoint response in *tdp1-AID wss1Δ*, while the additional *ulp1-ΔN* or *siz2Δ* mutations diminished its levels, as measured by the gel shift caused by Rad53 phosphorylation (Figure 2F). Moreover, both *siz2Δ* and to a lesser extend *ulp1-ΔN* rescued the G2/M-arrest observed in *tdp1-AID wss1Δ* cells upon auxin treatment (Figure 2G). Based on these observations, we surmise that sumoylation mutants promote more efficient Top1cc repair in *tdp1-AID wss1Δ*.

### Sumoylation changes in *ulp1-*Δ*N* and *siz2*Δ

Since elevated levels of Top1-DNA crosslinks underlie the *tdp1Δ wss1Δ* severe synthetic sickness (Stingele et al., 2014), and since Top1 is a known substrate for SUMO modification in budding yeast (Balakirev et al., 2015; Chen et al., 2007), we reasoned that it might modulate Top1cc processing. It was reported previously that Top1-SUMO levels are only weakly affected by single mutations in E3 SUMO ligases; in contrast, double *siz1 mms21* or *siz1 siz2*, but not *siz2 mms21* mutants strongly reduce Top1 SUMO conjugates (Chen et al., 2007). To evaluate the sumoylation status of Top1, we tagged the sole essential SUMO-encoding *SMT3* gene with 6His-Flag, allowing the isolation of sumoylated proteins by Ni-NTA beads. Contrary to the model that Top1 sumoylation accounts for the poor growth of *tdp1Δ wss1Δ* cells, the Top1-13Myc signal in the sumoylated protein fraction was decreased in *tdp1-AID wss1Δ* + auxin, as compared to wild-type (Figure 3A). Moreover, *ulp1-ΔN* and *siz2Δ* did not have the same effect on Top1 sumoylation: when combined with *tdp1-AID wss1Δ,* the *ulp1-ΔN* mutation increased Top1-SUMO levels, whereas *siz2Δ* had no appreciable effect (Figure 3A). To further evaluate a possible role of elevated Top1-SUMO levels in *ulp1-ΔN,* a *top1-3KR* mutant was created in which the three lysines previously implicated in bulk Top1 sumoylation were mutated to arginines (Chen et al., 2007). Whereas *top1-3KR* markedly decreased Top1 sumoylation (Figure S3A), this mutant neither had a deleterious effect on *ulp1-ΔN tdp1-AID wss1Δ,* nor improved the growth of *tdp1-AID wss1Δ* and its resistance to CPT (Figure 3B). Collectively, these results indicate that Top1 is not a Siz2 SUMO ligase target and argue against the model that Top1 sumoylation is responsible for the growth defect of *tdp1Δ wss1Δ*.

**Figure 3.**
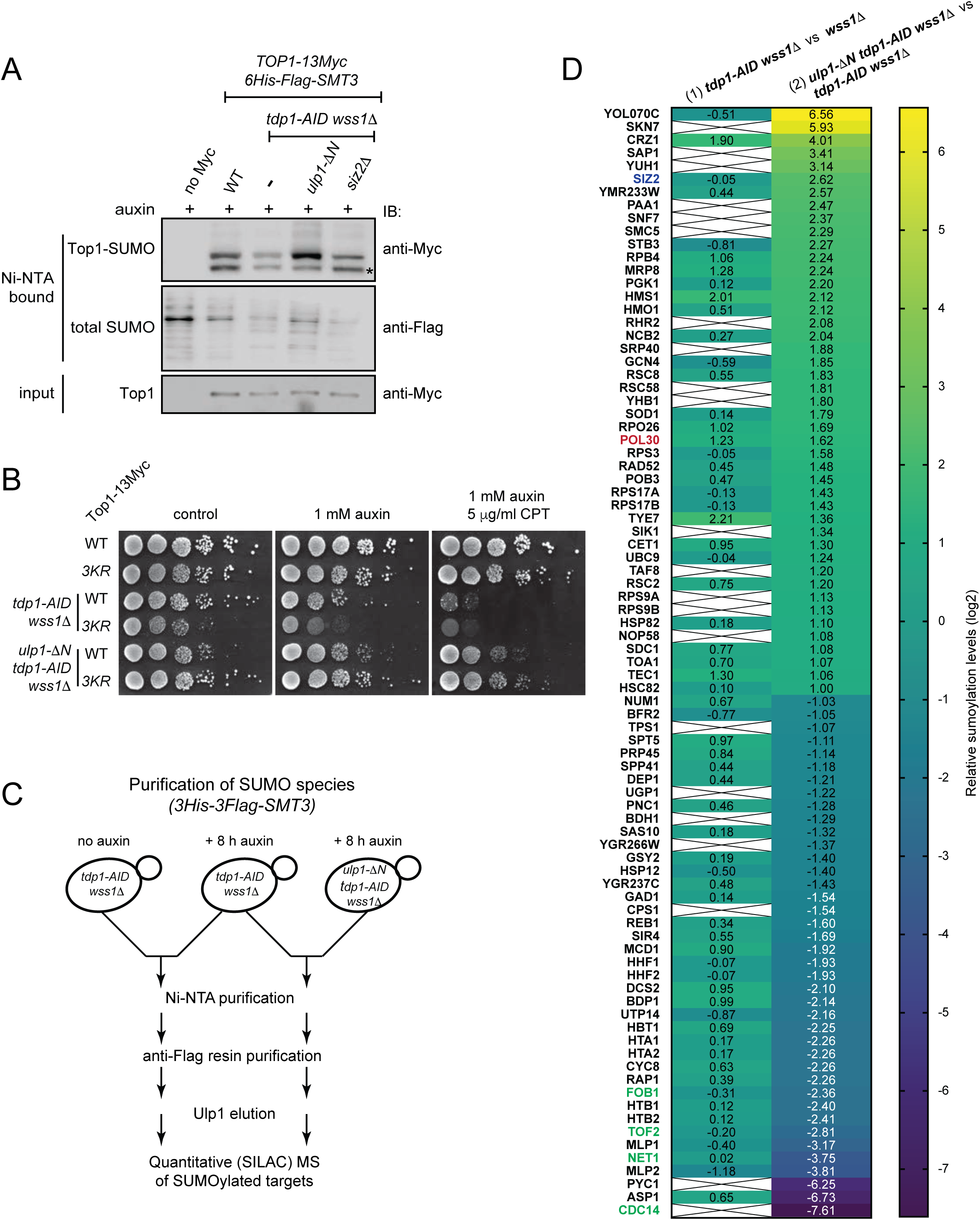
Top1 and global sumoylation changes in *ulp1-*Δ*N* and *siz2*Δ (A) Relative levels of sumoylated Top1 evaluated by the SUMO assay. *SMT3* (gene coding for yeast SUMO) and *TOP1* were tagged at genomic loci with 6His-Flag or 13Myc respectively on their C-terminus. Tdp1-AID*-6HA was depleted by 6 h auxin treatment. The asterisk indicates unmodified Top1-13Myc non-specifically bound to Ni-NTA beads. (B) Reduced Top1-SUMO levels do not promote suppression of *tdp1-AID wss1*Δ. *TOP1-13Myc* or *top1-3KR-13Myc* (K65R, K91R, K92R) replace the genomic copy of *TOP1*. Spot assays were plated on YEPD media and imaged 48h post-plating. See also Figure S3A for Top1-SUMO levels in different mutants. IB, immunoblotting. (C) Schematic representation of the quantitative SUMO proteomics technique. SILAC mass spectrometry was performed and analyzed as described in (Albuquerque et al., 2013). (D) Relative abundance of SUMO targets identified in two proteomes performed as shown in (C). Relative median ratios values (log2) are shown on the heatmap. At least 3 unique peptides were identified for each selected protein. Only proteins which changed sumoylation status >2 times in *ulp1-*Δ*N tdp1-AID wss1*Δ vs *tdp1-AID wss1*Δ are plotted. A cross indicates the absence or less than 3 unique peptides.

To identify candidate SUMO targets, we next applied quantitative SUMO proteomics, utilizing stable isotope labelling by amino acids in cell culture and mass spectrometry (SILAC-MS) (Albuquerque et al., 2013). Global sumoylation changes were examined in two sets of mutants: (1) *wss1Δ* vs *tdp1-AID wss1Δ*, and (2) *tdp1-AID wss1Δ* vs *ulp1-ΔN tdp1-AID wss1Δ*, as summarized in (Figure 3C). Proteins with a sumoylation change of at least 2-fold in the analysis (2) were used to generate a heat map (Figure 3D, log2 scale is used) and values from analysis (1) were added subsequently. All proteins identified in the two SUMO proteome analyses are listed in (Data S1).

Notably, the SUMO proteomics analysis provided a hint towards the mechanism of Siz1 and Siz2 degradation observed in *ulp1-ΔN* mutant (Figure 2B). In the proteomics experiment (Figure 3D, #2), the additional *ulp1-ΔN* mutation promoted 1.8-fold and 6-fold increase in sumoylated Siz1 and Siz2 respectively (Data S1 and Figure 3D). Hyper-sumoylation of Siz1 and Siz2 in *ulp1-ΔN* could be a signal for STUbL-dependent ubiquitination and degradation, as seen for their fission yeast ortholog Pli1 (Nie and Boddy, 2015). Consistent with this model, the budding yeast Slx5-Slx8 complex ubiquitinates Siz1 *in vivo* (Westerbeck et al, 2014) and conjugates ubiquitin to Siz2 *in vitro* (Mullen and Brill, 2008). Pli1 was proposed to be stabilized by Ulp1-mediated desumoylation, which is unlikely the case for Siz1 and Siz2, since the *ulp1-ΔN* mutation is dominant (Figure S1B).

We next selected from the SILAC-MS data a list of genes annotated with the ontology term “DNA repair” (Figure S3D). We were especially interested in the decrease of sumoylation in *ulp1-ΔN*, as these changes could have resulted from Siz2 degradation. Surprisingly, very few DNA repair factors had reduced SUMO levels (Figure S3D). Since DNA repair factors are among the least abundant cellular proteins (Ho et al., 2018), we cannot exclude that the current MS analyses lacked the sensitivity to identify more candidates from this group.

The *ulp1-ΔN* mutation also reduced sumoylation of several rDNA-related factors – Cdc14, Net1, Tof2, and Fob1 (Figure 3D, highlighted in green), consistent with the previous report that *ulp1-ΔN* is a gain-of-function mutant which desumoylates Ulp2 targets (de Albuquerque et al., 2016). Whereas we validated the decrease in Cdc14 sumoylation in the context of *ulp1-ΔN*, we did not observe a similar effect of *siz2Δ* (data not shown). We therefore decided not to further investigate the role of rDNA factors.

Sumoylation of the sliding DNA clamp PCNA, encoded by the *POL30* gene, was ∼3-fold higher in *ulp1-ΔN tdp1-AID wss1Δ* relative to *tdp1-AID wss1Δ* (Figures 3D and S3D). Posttranslational modifications of PCNA, including sumoylation, are known to regulate the choice between DNA damage repair or tolerance (Leung et al., 2018). To address the role of PCNA sumoylation in Top1cc processing, we analyzed the *pol30-2KR* mutant (lysines K127 and K164 are mutated to arginines), which lacks detectable sumoylation (Davies and Ulrich, 2012). A reduction in PCNA-SUMO species was observed in *tdp1-AID wss1Δ,* which was prominently reversed by *ulp1-ΔN* and *siz2Δ* (Figure S3B). However, the *pol30-2KR* mutant neither had an impact on the poor growth and CPT sensitivity of the *tdp1-AID wss1Δ*, nor on the *ulp1-ΔN-* and *siz2Δ*-mediated suppression (Figure S3C), suggesting that sumoylation of PCNA is not involved in this process.

### SUMO levels at the DPC site modulate the speed of DPC repair

Siz2 sumoylates multiple proteins involved in DNA replication and repair in response to certain genotoxin treatments (Cremona et al., 2012; Psakhye and Jentsch, 2012). If Top1cc also induces multiple sumoylation events at the damage site, the synthetic rescue effects of *ulp1-ΔN* and *siz2Δ* may not be attributable to a unique substrate. To test this idea, we used the Flp-nick system that generates a site-specific DPC bound to DNA through a chemical bond identical to Top1cc (Nielsen et al., 2009). In this system, galactose-induced Flp is targeted to the Flp recognition target (*FRT*) sequence, and the H305L mutation prevents the resolution of a recombination intermediate, leaving a covalent crosslink between Flp and DNA, or Flp-cc (Figure 4A). The *top1Δ* mutation allows to study Flp-nick defects independently of the Top1cc-related phenotypes. We have previously shown that *tdp1Δ wss1Δ top1Δ* cells are sensitive to Flp-cc induction, as reflected by the growth defect on galactose (Serbyn et al., 2020) (Figure 4B). Both *ulp1-ΔN* and *siz2Δ* mutations rescued this growth defect (Figure 4B), confirming that suppression by these SUMO pathway mutants is recapitulated in the Flp-nick system. Consistent with our hypothesis, chromatin immunoprecipitation targeting SUMO showed that Flp-cc induction was accompanied by a local increase in sumoylation as a function of proximity to the crosslinked *FRT* site (Figure 4C, compare raffinose and galactose). The *tdp1Δ wss1Δ top1Δ* mutant accumulated even higher SUMO levels in galactose, and the increase was only observed in the presence of the *FRT* binding site (Figure 4C). Remarkably, additional *ulp1-ΔN* and *siz2Δ* mutations significantly decreased the SUMO signal in the vicinity of *FRT* (Figure 4C). In contrast, this decrease was not observed in the additional *slx5Δ* mutant (a weak suppressor of *tdp1Δ wss1Δ*), suggesting that this STUbL functions downstream of the SUMO modification. Together, these observations imply that Siz2 places toxic sumoylation marks in the proximity of the DPC in the absence of Tdp1 and Wss1.

**Figure 4.**
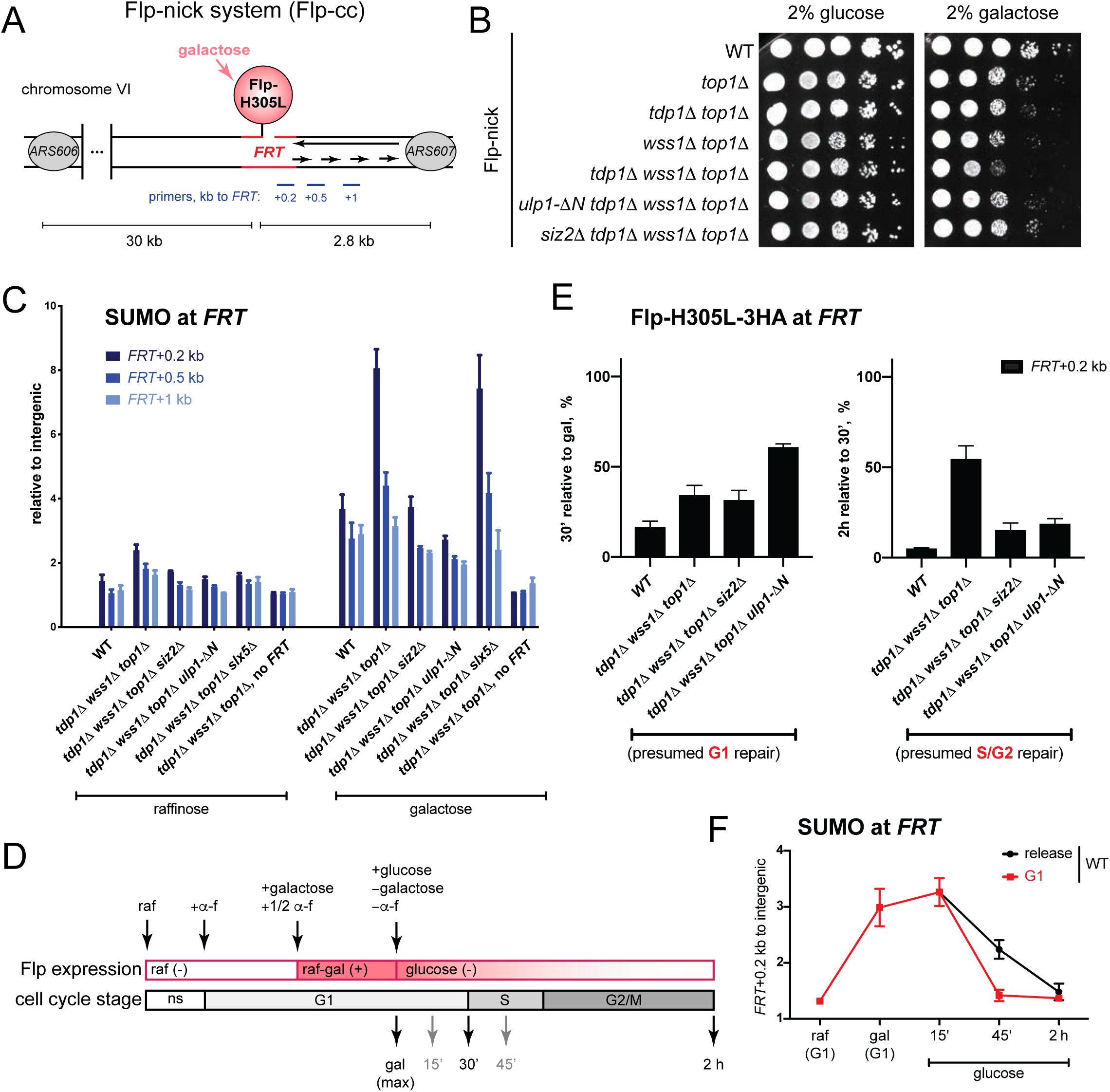
Excessive sumoylation by Siz2 in the vicinity of the DPC delays S-phase dependent repair of Flp-cc (A) Schematic representation of the Flp-nick system described in (Nielsen et al., 2009). Expression of the *pGAL10-flp-H305L* construct is induced by galactose; the mutant Flp recombinase is targeted to the *FRT* locus artificially introduced 2.7 kb upstream of the *ARS607* replication origin into chromosome VI; the system generates a DPC (Flp-cc) on the nicked DNA. Yeast cells are devoid of 2μ plasmid, a natural substrate of the Flp recombinase. (B) *ulp1-*Δ*N* and *siz2*Δ mutations rescue growth defects caused by Flp-nick induction in *tdp1Δ wss1Δ top1Δ*. Cells were pre-grown in YEP-2% raffinose medium prior to plating on SC- 2% glucose or SC- 2% galactose plates. Images were taken after 60 h of growth. (C) Sumoylation of the *FRT* locus. Non-synchronous yeast cultures were either grown in YEP+ 2% raffinose or additionally induced with 3% galactose for 2 h. Total SUMO (6His-Flag-Smt3) was immunoprecipitated with an anti-Flag antibody; ChIP-qPCR signals at the *FRT* locus (relative to intergenic) are plotted as the mean +/- SEM of 3-6 replicates. (D) Experimental design of Flp-nick induction in G1-synchronized cells. raf, raffinose; gal, galactose; *α*-f, alpha factor; ns, non-synchronous. (E) Kinetics of Flp-cc repair in different mutants. Yeast strains were grown as shown in (D). In addition to indicated mutations, all Flp-nick strains contain *bar1Δ* and *pGAL-flp-H305L-3HA.* Flp-H305L-3HA levels at *FRT* were monitored by ChIP-qPCR without crosslink using an anti-HA antibody. qPCR primers map 200 bp downstream of *FRT*. Data are presented as the mean +/- SEM of 5 independent replicates. The percent of input for each genotype was normalized to maximum galactose induction (left) or to the 30’ release timepoint (right). See also S4A for the cell cycle analysis of different mutants and S4D for glucose release in G1 and the normalization of all timepoints to galactose. (F) SUMO dynamics at *FRT* in different cell cycle stages. Cells were grown as shown on Figure S4B and further used for SUMO ChIPs with anti-Flag antibody. ChIP signals are presented as the mean±SEM fold increase of *FRT*+0.2kb over the intergenic region for n=4.

If the excessive sumoylation at the *FRT* locus is inhibitory for repair, *ulp1-ΔN* or *siz2Δ* should accelerate the kinetics of Flp-cc turnover. To test this possibility, cells were synchronized by *α*-factor in G1, *pGAL10-flp-H305L-3HA* expression was induced by galactose, followed by the simultaneous glucose repression and release into the cell cycle (Figure 4D). Immunoprecipitation was used to assess the levels of Flp-H305L-3HA crosslinked to *FRT* in galactose (maximum induction in G1), after 30 min glucose release (corresponding to G1/early S), or 2 h glucose release (G2) (Figures 4D and S4A). Because starting Flp-cc levels in galactose considerably differed between the mutants, the results were presented as % of remaining signal relative to the previous timepoint. As reported before (Serbyn et al., 2020), *tdp1Δ wss1Δ top1Δ* significantly slows down the speed of Flp-cc repair. After 30 min glucose release, Flp-H305L-3HA remained crosslinked to DNA in ∼16% of WT-like cells versus ∼34% in *tdp1Δ wss1Δ top1Δ* (Figure 4E, left). The additional *siz2Δ* mutation had a similar to *tdp1Δ wss1Δ top1Δ* repair kinetics, and *ulp1-ΔN* counterintuitively further slowed down Flp-cc repair (Figure 4E, left). The 30 min timepoint corresponds to G1 or early S cell cycle stage (Figure S4A), suggesting that sumoylation does not inhibit repair in G1. Strikingly, however, between 30 min and 2 h, >50% of *tdp1Δ wss1Δ top1Δ* cells failed to repair the remaining Flp-cc, while this number dropped to only 15-20% in the cells containing additional *siz2Δ* or *ulp1-ΔN* mutations (Figure 4E, right). From 30 min to 2 h glucose release, Flp-cc repair presumably takes place in the context of S/G2, suggesting that excessive sumoylation is deleterious for cells progressing through these cell cycle phases.

We next tested how the cell cycle arrest in G1 impacts the kinetics of Flp-cc repair in different mutants. As before, Flp-cc was induced in *α*-factor-synchronized cells, but this time cells were retained in G1 after shift to glucose (Figures S4A and S4B). Of note, wild-type cells were proficient for Flp-cc repair even in the absence of replication, and *tdp1Δ wss1Δ top1Δ* repaired more slowly (Figure S4C). In line with the slower Flp-cc processing after 30 min release (Figure 4E, left), the *ulp1-ΔN tdp1Δ wss1Δ top1Δ* was deficient for the G1-dependent repair even after 2 h of *α*-factor arrest (Figure S4D). The *siz2Δ tdp1Δ wss1Δ top1Δ* mutant did not show as strong a defect, however it still processed Flp-cc more slowly than WT-like cells in G1 (Figure S4D). Thus, acceleration of repair by the *siz2Δ* and *ulp1-ΔN* mutations requires progression to S and/or G2 phases.

Since Flp-cc formation promotes a local increase in sumoylation (Figure 4C), we addressed next how cell cycle stage impacts SUMO dynamics at *FRT*. Remarkably, SUMO species were turned over more efficiently if cells were retained in the G1 phase (Figure 4F). The same trend was observed in the *tdp1Δ wss1Δ top1Δ* mutant (Figures S4E and S4F). Together, the above results suggest that sumoylation promotes DPC processing in G1, but that the remaining excessive SUMO species delay the replication-dependent repair in the absence of Tdp1 and Wss1. This delay can be suppressed at least in part by lowering sumoylation via the *siz2Δ* or *ulp1-ΔN* mutations.

### Alternative Top1cc repair pathways function in *tdp1*Δ *wss1*Δ *siz2*Δ

Ulp1- and Siz2-dependent suppression of *tdp1Δ wss1Δ* implies that the loss of sumoylation activates alternative repair pathways that deal with Top1ccs. To identify those pathways, a SATAY transposon screen was performed on *tdp1Δ wss1Δ siz2Δ*, as shown on Figure 5A. The transposon coverage in the mutant *tdp1Δ wss1Δ siz2Δ* SATAY library was compared to a pool of several libraries wild type for these genes (Figure 5B) and (Data S2). As *TDP1, WSS1,* and *SIZ2* were deleted and therefore could not be targeted by transposons, these genes appeared as top hits in the screen. Potential synthetic lethal genetic interactors of *tdp1Δ wss1Δ siz2Δ* are highlighted on the volcano plot (Figure 5B). Among them, several known Top1ccs- and DNA double strand break repair factors were found, which are analyzed next.

**Figure 5.**
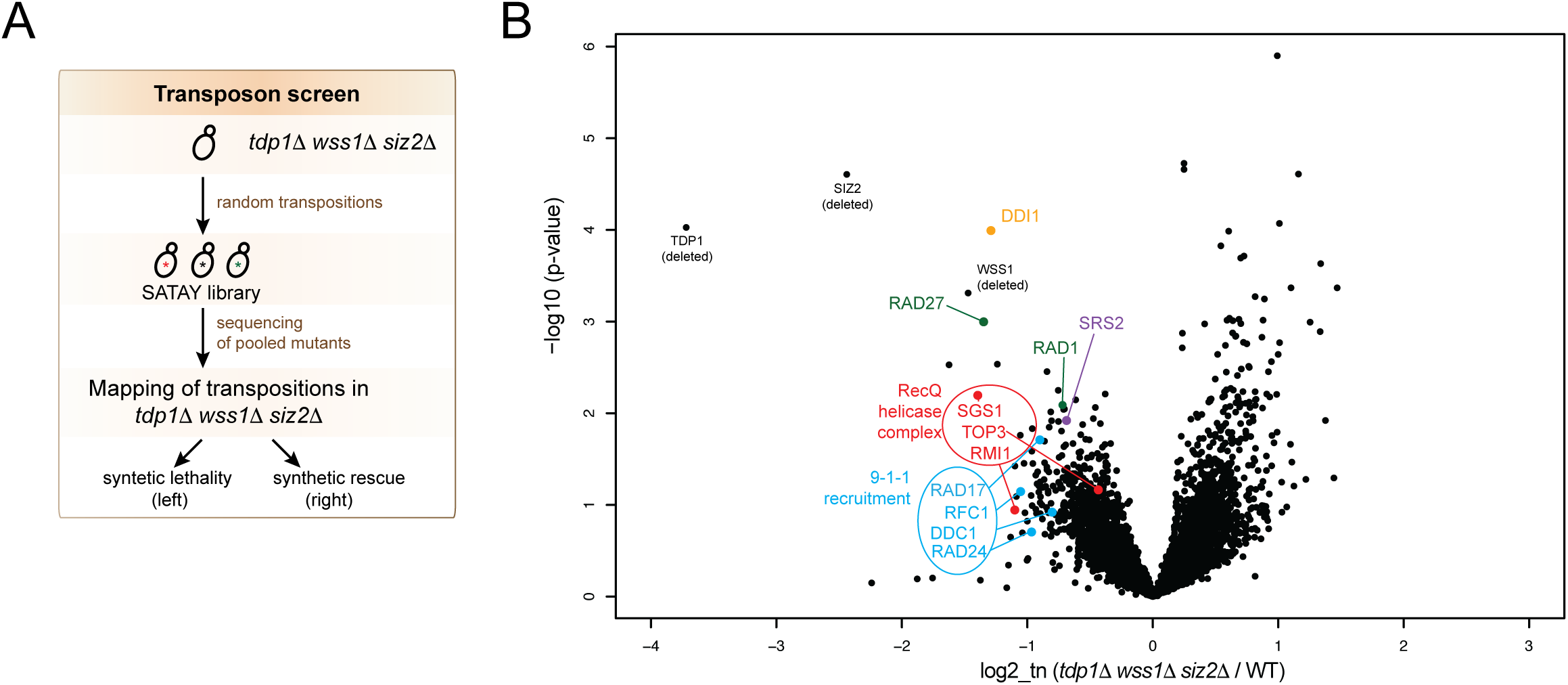
A genetic screen uncovers factors essential for *tdp1*Δ *wss1*Δ *siz2*Δ survival (A) Outline of the SATAY transposon screen designed to identify genes required for *tdp1*Δ *wss1*Δ *siz2*Δ survival. (B) A volcano plot summarizing results of the SATAY transposon screen of *tdp1*Δ *wss1*Δ *siz2*Δ performed as described in (A). The fold-change in the number of transposons per yeast gene body in *tdp1*Δ *wss1*Δ *siz2*Δ library obtained by comparison to a pool of six unrelated libraries (log2, x-axis). Respective p-values (-log10, y-axis) are plotted. Selected DNA damage repair factors are highlighted in colors. For validations and additional genetic analyses, see Figure S5, Tables S1 and S2.

Several structure-specific endonucleases are thought to eliminate Top1ccs from DNA in the absence of the direct Top1-DNA bond cleavage by Tdp1 (Deng et al., 2005; Hartsuiker et al., 2009; Liu et al., 2002; Reid et al., 2011; Vance and Wilson, 2002; Zheng et al., 2005). Consistently, the genes encoding the Rad27 and Rad1 nucleases were among the least targeted in the *tdp1Δ wss1Δ siz2Δ* SATAY screen (Figure 5B). To further test the importance of structure-specific nucleases, we deleted *RAD27, MRE11* or disrupted several other nuclease complexes found in the screen or previously implicated in Top1cc processing – Rad1-Rad10, Slx4-Slx1, and Mus81-Mms4. Most of these mutations modestly affected growth and CPT resistance of the *tdp1Δ wss1Δ siz2Δ* strain (summarized in Tables S1 and S2). In contrast, *RAD27*, a gene that encodes a flap nuclease with a dual role in DNA replication and DNA damage repair, was essential for *tdp1Δ wss1Δ siz2Δ* viability (Figure S5A). Combination of individual or double *tdp1Δ*, *wss1Δ*, or *siz2Δ* mutations with *rad27Δ* was not detrimental for cell growth. These data suggest that Rad27 is indispensable for Top1cc repair when Tdp1-, Wss1-, and Siz2-dependent pathways are not accessible.

DNA helicases are known to play important roles in diverse DNA repair pathways, including the removal of DPCs. A number of them were identified in the SATAY screen (Figure 5B). The ReqQ helicase Sgs1 (BLM and WRN in mammals) works with Top3 to resolve Holliday junctions and to promote the restart of replication forks collapsed at Top1ccs (Pichierri et al., 2000; Vance and Wilson, 2002). The *sgs1Δ* mutation was previously reported to be deleterious in *wss1Δ* (Mullen et al., 2010); loss of this helicase in *tdp1Δ wss1Δ siz2Δ* did not cause additional growth defects, as compared to *sgs1Δ wss1Δ* (Figure S5B). Another helicase, the Srs2 anti-recombinase, is recruited to the site of damage *via* interactions with sumoylated PCNA (Pfander et al., 2005) and disrupts Rad51 nucleoprotein filaments formation (Krejci et al., 2003). Interestingly, the combination of *srs2Δ* and *siz2Δ* alone was deleterious in the presence of the Top1cc-inducing drug CPT, a phenotype reflected also in the context of *tdp1Δ wss1Δ siz2Δ* (Figure S5C). These findings suggested an involvement of these helicases in processing Top1ccs in *tdp1Δ wss1Δ siz2Δ*, confirming the findings from the SATAY screen. The DNA damage checkpoint was shown to play a role in the response to CPT-induced

DNA damage (Pommier et al., 2006; Simon et al., 2000). Consistent with this idea, the 9-1-1 complex (Ddc1, Rad17, Mec3) and its loader Rad24 were all identified in the SATAY screen (Figure 5B). Further genetic analysis showed that the *rad17Δ* and *rad9Δ* checkpoint response mutants both sensitize *tdp1Δ wss1Δ siz2Δ* to CPT, although they did not appreciably affect the growth property of this triple mutant (Figures S5D and S5E).

The Ddi1 protease previously implicated in DPC processing (Serbyn et al., 2020) was also found as a top candidate in the screen (Figure 5B). Ddi1 was required for the CPT resistance of *tdp1Δ wss1Δ siz2Δ* (Figure S5F), further supporting that Ddi1 may function as a backup Top1cc proteolysis pathway.

Taken together, multiple DNA repair and checkpoint pathways play a role in improving the growth of *tdp1Δ wss1Δ siz2Δ* and its response to CPT-induced DPCs. They include endonucleases, helicases, the Ddi1 DPC protease, as well as the DNA damage checkpoint genes. Thus, reducing sumoylation at DPCs in the *tdp1Δ wss1Δ* mutant appears to trigger the activation of an extensive network of DNA repair and DNA damage signalling pathways that collectively counter the toxic effect of DPCs.

## DISCUSSION

DPC adducts, including Top1ccs, are formed during normal cell metabolism but do not threaten cell survival, owing to multiple DPC repair mechanisms involved in their elimination. The redundancy in such DNA repair pathways becomes especially important when cells are exposed to diverse industrial, household and environmental agents or chemotherapeutic drugs that induce DPCs. In this study we show that SUMO helps to create a balance between the redundant Top1cc repair pathways in budding yeast. We reveal that the E3 ligase Siz2 conjugates SUMO in the proximity of a damaged site. Our genetic findings suggest that SUMO stimulates repair by promoting the recruitment of the DPC protease Wss1. At the same time, we show that excessive Siz2-dependent SUMO conjugation inhibits alternative repair mechanisms, and additionally identified at least some of them in a genetic screen. Moreover, excessive sumoylation of the unrepaired DPC by Siz2 is especially deleterious outside of the G1 cell cycle phase. How SUMO could cooperate with some or compete with other DPC repair mechanisms to coordinate them in time and space is discussed below.

### Favorable effects of SUMO conjugation around the DPC locus

The cellular response to DPC-induced DNA damage can be divided into several steps: (1) DPC detection, (2) proteolysis and (3) elimination of the remaining DNA damage (Figure 6). The SUMO signal increases in the vicinity of the DPC-containing locus upon damage formation (Figure 4C) and likely facilitates DPC detection (Figure 6, step 1). Conjugation of SUMO was previously documented in several DPC models such as TOP1, TOP2, M.HpaII, and DNMT1 (Borgermann et al., 2019; Larsen et al., 2019; Mao et al., 2000a; Mao et al., 2000b; Schellenberg et al., 2017), suggesting a general response mechanism. SUMO could both mark the covalently trapped protein directly and/or be conjugated to the other factors residing in close proximity to the DPC. This was observed for the Top1cc and Flp-cc models used in this study (Figures 3A and 4C) (Chen et al., 2007; Xiong et al., 2009), and seen previously for the trapped mammalian DNMT1 (Borgermann et al., 2019). Our data allow to distinguish between the two events and suggest that SUMO conjugation by Siz2 around the DPC, but not to Top1, is detrimental for Top1cc repair. One interpretation of these data is that multiple SUMO conjugation events around a DPC create a sumoylated domain required to recruit relevant factors and establish a platform for efficient DNA repair. In line with this model, Siz2 sumoylates numerous DNA repair proteins in response to other genotoxin treatments such as MMS or zeocin (Chung and Zhao, 2015; Psakhye and Jentsch, 2012). Consistently, our SILAC-MS analysis and the subsequent candidate approach does not point to a single Siz2 target relevant for DPC repair (Figures 3 and S3).

**Figure 6.**
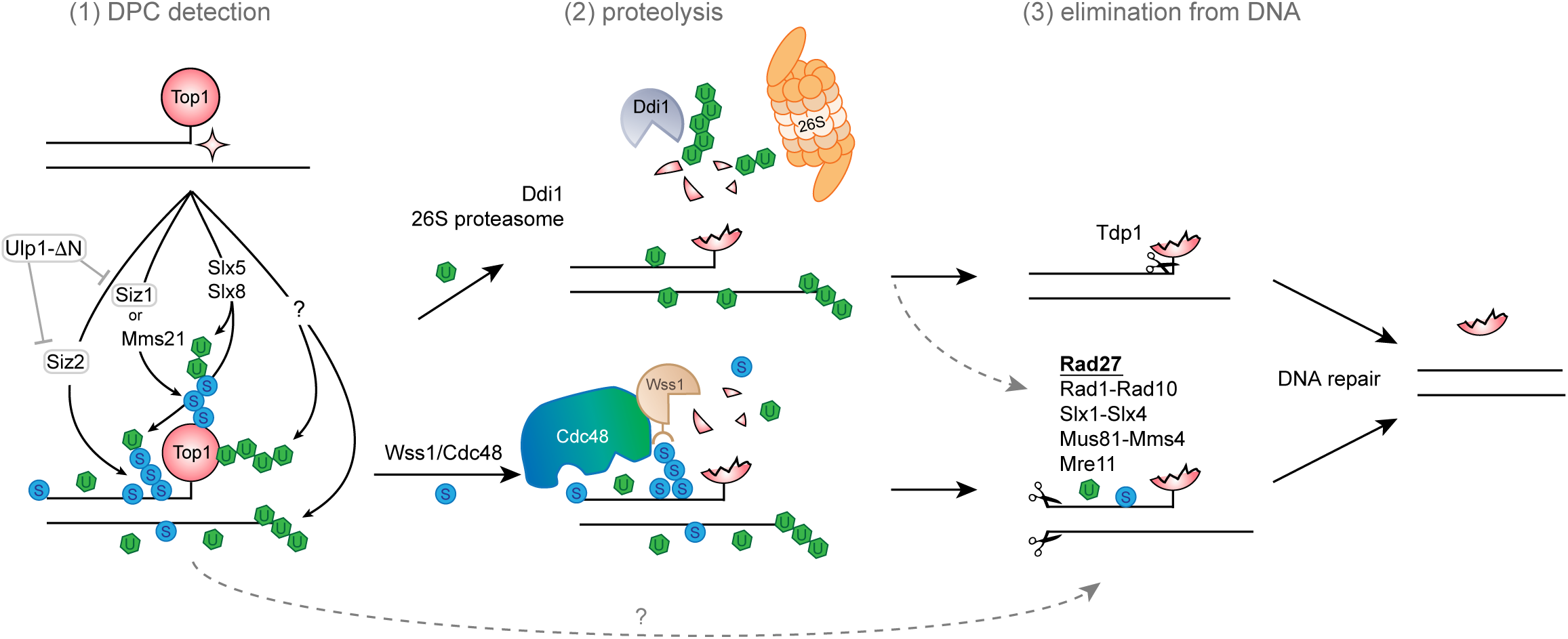
A hypothetical model summarizing the interplay between PTMs and different DPC repair factors Topoisomerase 1 (Top1) is covalently trapped on DNA as a result of an aberrant enzymatic reaction, forming a DPC referred to as Top1cc. Top1cc and proximally located proteins are modified by SUMO (S) and ubiquitin (U). (1) E3 SUMO ligases Siz1 and Mms21 facilitate Top1 sumoylation; Siz2 sumoylates proteins at the DPC lesion site. DPCs are additionally ubiquitinated in a SUMO-dependent manner (by the Slx5-Slx8 complex) or independently of SUMO (by unknown E3 ligase(s) in yeast). (2) SUMO stimulates Wss1 recruitment; ubiquitin and mixed SUMO-Ub chains are recognized by Cdc48 (Wss1 partner); ubiquitin is also recognized by the alternative DPC proteases Ddi1 and 26S proteasome. Either of these three proteases allows the initial degradation of a protein trapped in DPC. (3) Partial or full Top1cc proteolysis opens access for Tdp1 cleavage (preferred pathway) or allows DNA cleavage by various structural endonucleases, such as Rad27, Rad1-Rad10, Slx1-Slx4, Mus81-Mms4, and Mre11. SUMO and ubiquitin may additionally influence the downstream DPC repair. Excessive sumoylation that persists in the absence of Wss1 and Tdp1 (presumably G1-dependent) will inhibit efficient Top1cc processing in the S phase of the cell cycle (not shown).

In line with this model, loss of SUMO acceptor lysines in Top1 has no or only a minor effect on Top1cc repair in *tdp1Δ wss1Δ* (Figure 3B). The *top1-3KR* mutant was described previously to abrogate most of Top1 sumoylation (Chen et al., 2007), yet it produces no phenotype in the context of the tested mutants (Figure 3B). Of note, another recent study found a partial suppression of *tdp1Δ wss1Δ* by *top1-4KR* (Sharma et al., 2017). The *top1-4KR* mutant has an additional K600R amino acid substitution, but this mutation does not further reduce Top1 sumoylation, as compared to *top1-3KR* (Chen et al., 2007). The K600R mutation is therefore unlikely at the basis of the discrepancy with our results. Moreover, high molecular weight sumoylated species of Top1 remain detectable in both *top1-3KR* and *top1-4KR* mutants (Figure S3A) (Chen et al., 2007), raising the possibility that additional sumoylation of Top1 can still be involved. A comprehensive identification of Top1 sumoylation sites might provide a definitive answer about the role of Top1 sumoylation in Top1cc removal.

What factor could SUMO recruit to the DPC locus? The Wss1 DPC protease is an obvious candidate as it has strong SUMO binding affinity indispensable for its *in vivo* proteolytic function in yeast (Balakirev et al., 2015; Mullen et al., 2010; Stingele et al., 2014). Cdc48, a Wss1 partner essential for Top1cc repair, recognizes mixed SUMO-Ub signals through an adaptor protein (Nie et al., 2012); it could therefore also stimulate Wss1 recruitment to a sumoylated locus. Although we did not directly test Wss1 recruitment here, our genetic data suggest that Wss1 and the E3 SUMO ligase Siz2 function in the same pathway (Figure 2E). All these observations collectively suggest that the main function of SUMO at the DPC locus is likely to promote Wss1 recruitment directly or via Cdc48 (Figure 6). Among Wss1 orthologues in higher eukaryotes, GCNA also has a SUMO binding motif (Borgermann et al., 2019; Dokshin et al., 2020), indicating that DPC control by SUMO could be evolutionarily conserved.

The Slx5-Slx8 STUbL is another complex that could be directed by SUMO to DPC sites. STUbLs are described to facilitate Top1cc processing in budding, fission yeast, and mammalian cells (Nie et al., 2017; Sharma et al., 2017; Sun et al., 2019). The Slx5-Slx8 STUbL promotes repair in a pathway parallel to Tdp1 (Figure S1F), as also observed in fission yeast (Nie et al., 2017). Mixed SUMO-Ub chains generated by the Slx5-Slx8 complex could, on the one hand, recruit Cdc48-Wss1 (Figure 6), as suggested by the epistatic relationship of *wss1Δ* with the *tdp1Δ slx5Δ* mutant (Figure S1F). On the other hand, ubiquitin could stimulate Top1cc degradation by 26S proteasome (Figure 6), as proposed for the mammalian Slx5-Slx8 functional orthologue RNF4 (Sun et al., 2019).

### Deleterious effects of Siz2-dependent SUMO conjugation on Top1cc processing

Wss1 is a low abundance protein in yeast (Stingele et al., 2015); its availability could therefore be rate-limiting for DPC repair. When cells are challenged with an excess of DPCs, they may require an alternative DPC protease. Ddi1 and 26S proteasome could at least partially substitute for Wss1 function (Serbyn et al., 2020), and both of them target ubiquitinated proteins for degradation (Finley et al., 2012; Yip et al., 2020). Ubiquitin and SUMO are known to compete for the acceptor lysines (Hendriks et al., 2014). Thus, excessive sumoylation by Siz2 could block the access of alternative proteases to the lesion in *tdp1Δ wss1Δ* (Figures 4C and 2A). In agreement with this model, Ddi1 appears especially important when excessive sumoylation in *tdp1Δ wss1Δ* is prevented by the *siz2Δ* mutation (Figures 5B and S5F).

At least some repair pathways identified in the *tdp1Δ wss1Δ siz2Δ* transposon screen could assist Ddi1 or 26S proteasome in downstream damage processing (Figure 6). Structure-specific nucleases may be required to clear the peptide remnants, especially when Tdp1 is absent or not accessible and fails to perform direct digestion of the Top1-DNA bond. The identified nucleases and helicases are known to recognize and resolve aberrant replication and recombination intermediates that appear during or post-DNA replication, consistent with our earlier observations that Ddi1 signal appears at the DPC locus in a replication-dependent manner (Serbyn et al., 2020). Moreover, almost every factor described above is a SUMO target itself (Sarangi and Zhao, 2015). Molecular effects of sumoylation greatly vary depending on the substrate and sometimes could be inhibitory for repair; we cannot exclude that excessive SUMO additionally hampers one or several downstream DPC repair steps.

### Spatial and temporal control of DPC sumoylation

If several mechanisms compete for DPC repair, cells may have separated them in time and/or space. One possibility is that DPCs could be differently processed depending on the cell cycle stage. For example, in a *Xenopus* cell-free system, the ubiquitination by TRAIP is one of the earliest events accompanying replication-coupled DPC resolution, whereas the same DPC undergoes extensive sumoylation in the absence of replication (Larsen et al., 2019). Consistently, we observe DPC sumoylation in G1 (Figure 4C), and our data suggest that SUMO becomes inhibitory for the S-phase dependent DPC processing, at least in the context of *tdp1Δ wss1Δ* (Figure 4E). In wild-type cells, SUMO could help to cope with low or moderate DPC levels in G1. In *tdp1Δ wss1Δ* or in similar situations when DPC levels are elevated, an overwhelming amount of such lesions will be carried to S-phase. In this scenario, SUMO attached to the non-processed DPCs would inhibit the replication-dependent repair.

Intranuclear positioning is known to determine the fate of diverse damaged loci, including heterochromatic DSBs, collapsed replication forks, eroded telomeres, and repeated DNA sequences. SUMO stimulates the motion of the respective loci to the nuclear periphery to define the succeeding DNA repair pathway choice (Amaral et al., 2017; Geli and Lisby, 2015). We and others showed that proper NPC anchoring of the Ulp1 SUMO protease impacts Top1cc processing (Figures 1 and 2C) (Chen et al., 2007), suggesting that DPC positioning inside the nucleus may also be detrimental for damage processing. It remains to be determined whether hyper-sumoylation of DPCs could be directly reverted at the NPC by Ulp1 and how such an event may affect Top1cc processing.

In conclusion, the SUMO posttranslational modification tunes available alternative DPC repair pathways. Substantial additional work is required to understand all mechanistic details and to identify the exact sumoylation event(s) capable to alter DPC processing. Given the evolutionary conservation of mechanisms involving the SUMO modification in DNA damage response, these new aspects of DPC biology are anticipated to be relevant for different domains of life. Last, the suppressive phenotypes of SUMO pathway mutants could help to understand the resistance to Top1 inhibitors often observed in chemotherapy.

## Supporting information

Table S3 - strains_plasmids_oligonucleotides_mutations

Data S1 - SUMO proteomics

Data S2 - tdp1wss1siz2 SATAY

## ACKNOWLEDGEMENTS

We thank Helle Ulrich for kindly sharing the parental auxin degron strain, PCNA plasmid constructs, anti-PCNA, and anti-SUMO antibodies; Lotte Bjergbaek for the Flp-nick system; Mark Hochstrasser for Ulp1 constructs; Benoît Palancade for *siz1Δ*, *siz2Δ*, and *mms21-11* strains; Takeo Usui for the *12geneΔ0HSR* strain. We are grateful for Geraldine Silvano’s technical assistance. We thank Maksym Shyian, Simon Boulton, Johannes Walter, Vincent Dion, Daria Gudkova, and all members of the Stutz laboratory for their comments, discussions, suggestions and critical reading of the manuscript. This work was supported by funds from the Swiss National Science Foundation (grants 31003A 153331 and 31003A_182344 to FS) and the Canton of Geneva. The Kornmann lab is supported by grants from the Swiss National Science Foundation (31003A_179549) and the Wellcome Trust (214291/Z/18/Z). R.T.S. was supported by a Postdoctoral Fellowship from the National Cancer Institute (NCI T32 CA009523). H.Z. was supported by funds from NIH R01 GM116897 and NIH S10 OD023498.

## AUTHOR CONTRIBUTIONS

Conceptualization: N.S., F.S. Methodology: N.S., I.B., A.H.M., R.T.S., H.Z., B.K., F.S. Investigation: N.S., I.B., A.H.M., R.T.S. Data Curation: N.S., B.K., A.H.M., R.S. Writing – Original Draft: N.S. Writing – Review and Editing: N.S., F.S., B.K, A.M, H.Z. Supervision: F.S., B.K., H.Z. Funding acquisition: F.S.

## Supplemental Information

**Supplemental Figure 1.**
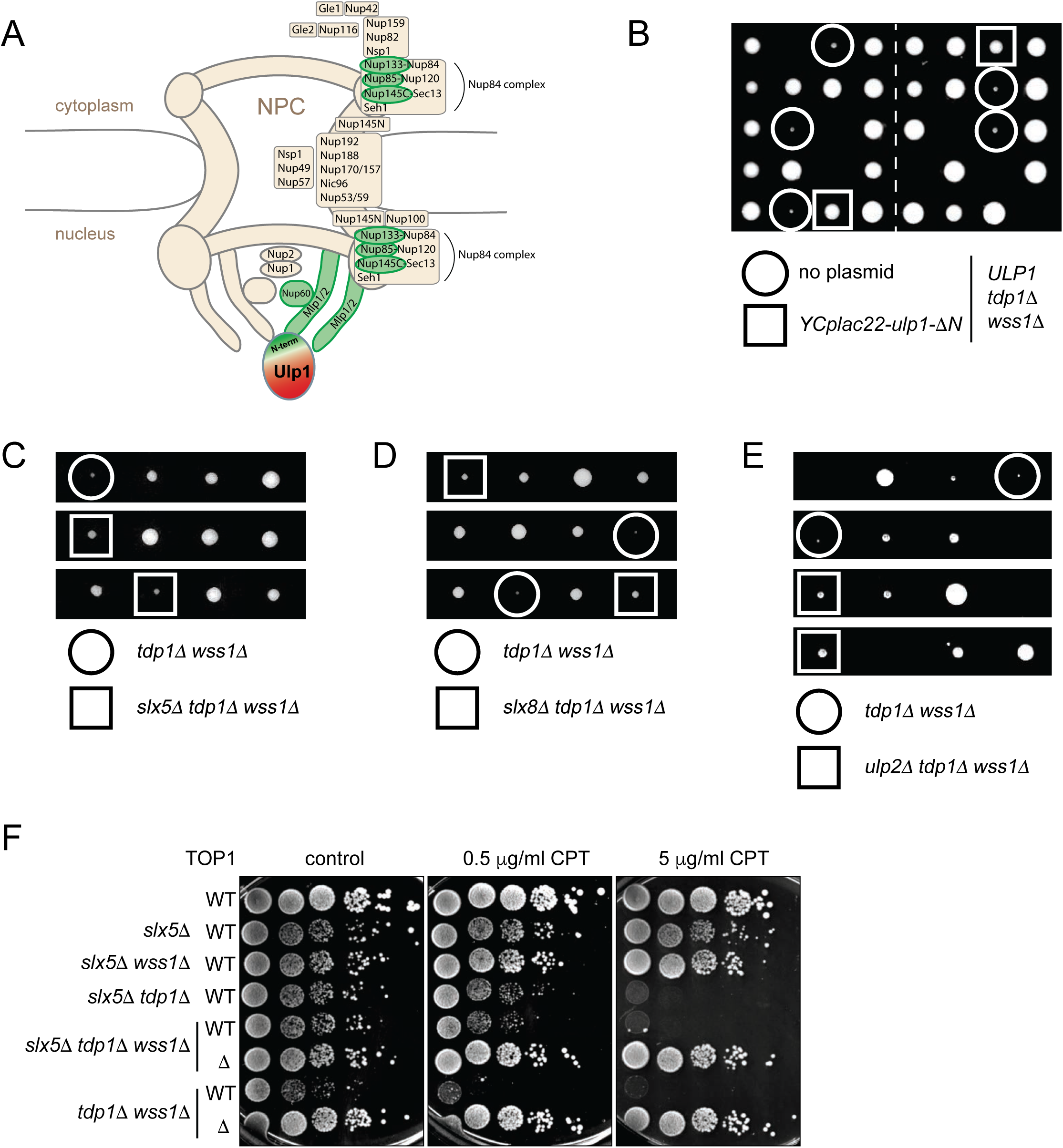
(related to Figure 1) (A) Nuclear pore proteins identified in the *tdp1-AID wss1*Δ + auxin SATAY genetic screen are present at the nuclear side of the NPC, where they anchor the Ulp1 SUMO protease to the nuclear periphery. Schematic representation of the NPC is based on (Beck and Hurt, 2017). Nucleoporins identified in the screen as suppressors of the *tdp1-AID wss1*Δ + auxin are marked in green. (B) The suppression effect of *ulp1-*Δ*N* is dominant. Tetrad analysis of the [*tdp1Δ/TDP1; wss1Δ/WSS1*] diploid transformed with the YCPlac22 centromeric plasmid expressing *ulp1-*Δ*N* from the *ULP1* promoter. (C-E) Loss of Slx5 and Slx8 SUMO-dependent ubiquitin ligases or Ulp2 SUMO protease weakly suppresses the growth defect of *tdp1*Δ *wss1*Δ. Tetrad analysis of [*tdp1Δ/TDP1; wss1Δ/WSS1*] diploids combined with *slx5Δ/SLX5* (C), *slx8Δ/SLX8* (D), and *ulp2Δ/ULP2* (E). (G) Loss of Slx5 sensitizes *tdp1*Δ to CPT but rescues the growth defect of *tdp1*Δ *wss1*Δ. 10x serial dilutions of yeast cultures were grown on YEPD plates in the presence of indicated amounts of CPT and imaged 48 h post-plating.

**Supplemental Figure 2.**
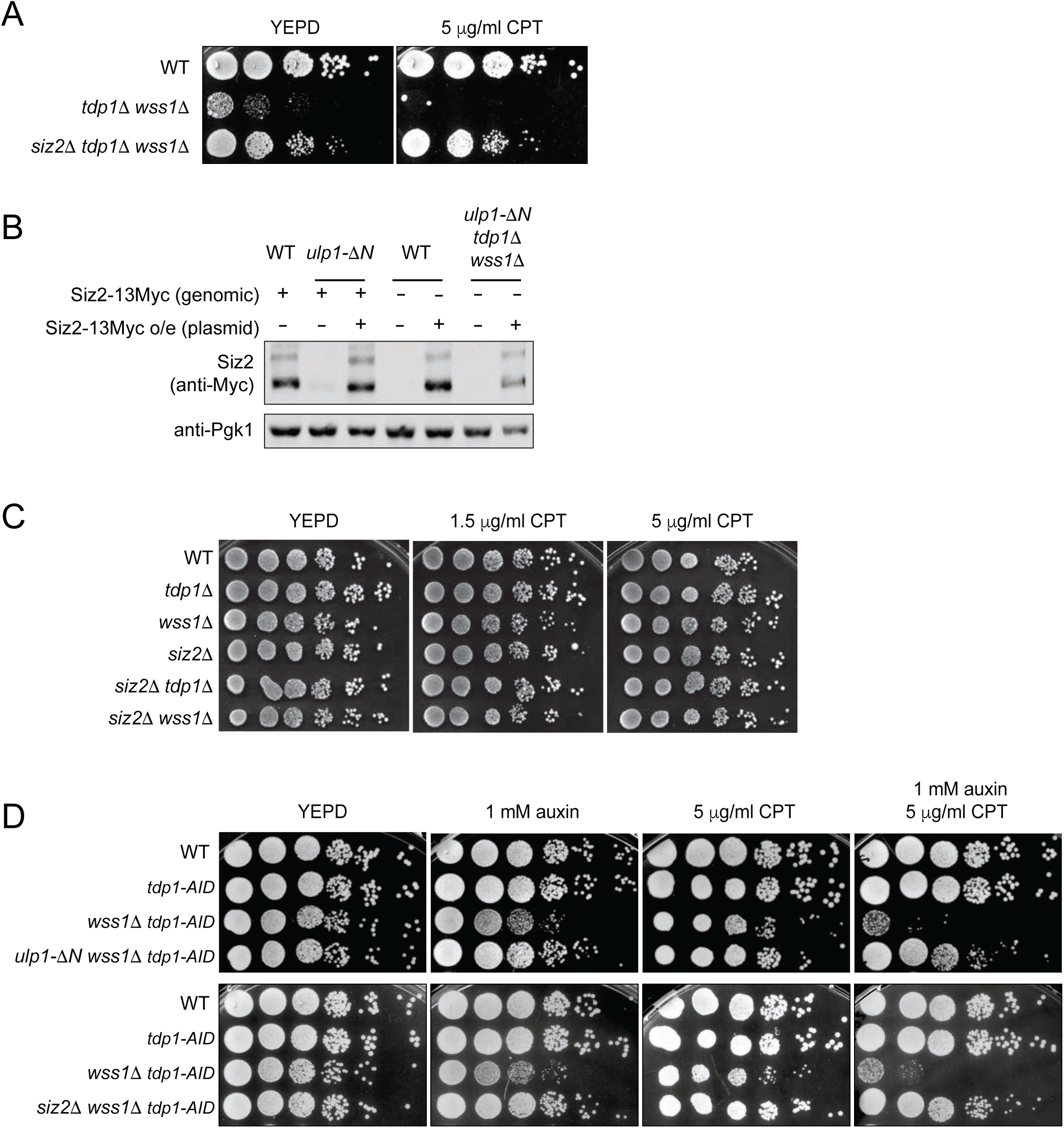
(related to Figure 2) (A) Absence of Siz2 efficiently suppresses CPT sensitivity of *tdp1*Δ *wss1*Δ. Indicated yeast strains were grown on YEPD plates in the presence or absence of CPT and imaged after 48 hours. (B) Siz2 protein levels are restored to WT-like levels by Siz2 overexpression from a plasmid carrying a strong *ADH* promoter (*p415-pADH-Siz2-13Myc-LEU2*). Control strains were transformed with an empty p415 plasmid. Where indicated, the genomic version of *SIZ2* was fused at the C-terminus with a 13Myc tag. Cell lysates from indicated mutants were subjected to immunoblotting with anti-Myc antibody. (C) Single and double *tdp1*Δ, *wss1*Δ, and *siz2*Δ mutants in the W303 genetic background are not sensitive to CPT. Spot test was performed as in (A). (D) Transient degradation of Tdp1 by the auxin degron system (*tdp1-AID*) mimics *tdp1*Δ phenotype in the presence of auxin. Spot assays were imaged after 48 h of growth.

**Supplemental Figure 3.**
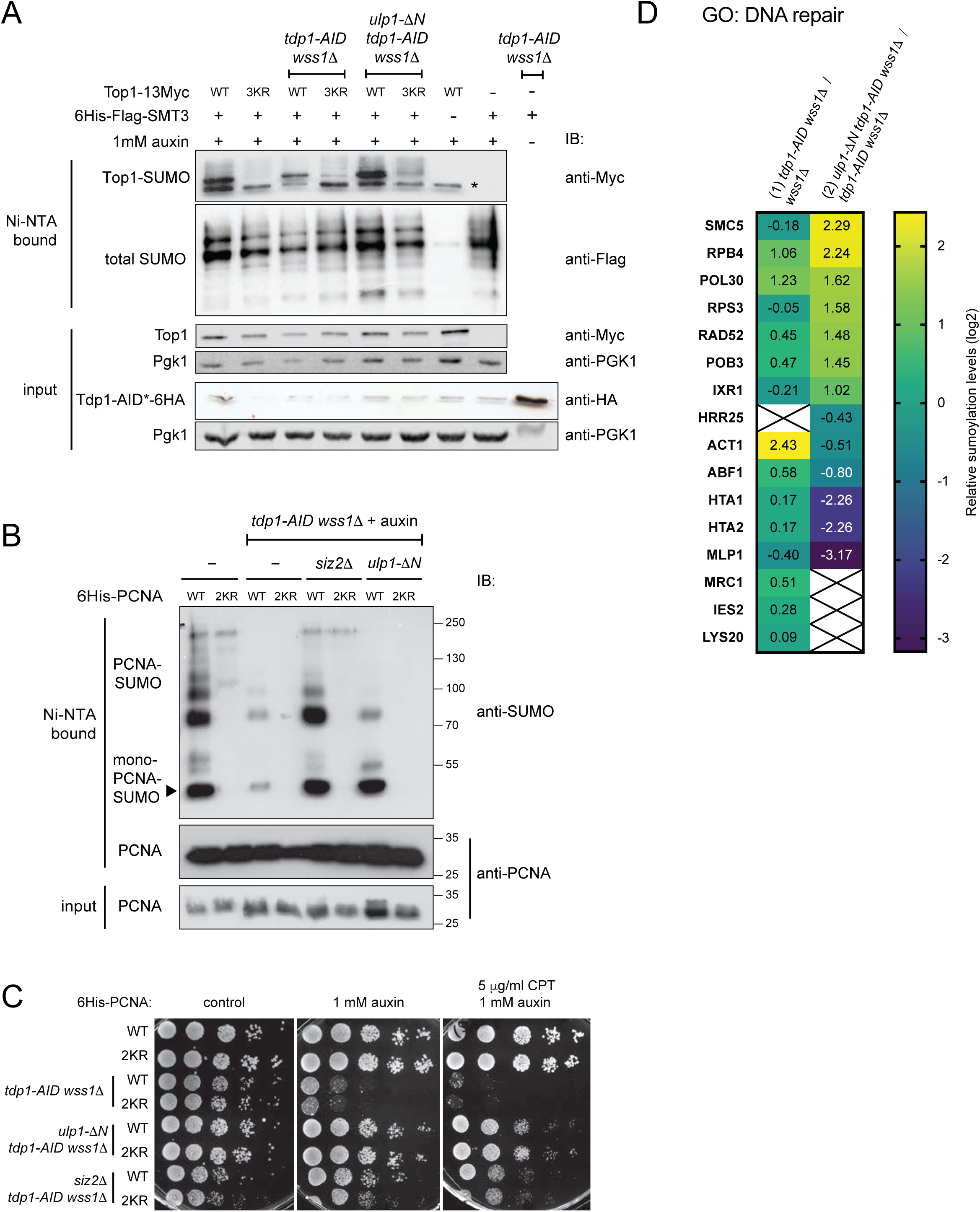
(related to Figure 3) (A) Sumoylation status of Top1-13Myc or Top1-3KR-13Myc mutants shown on Figure 3B. The SUMO assay was performed as in Figure 3A. IB, immunoblotting. (B) PCNA sumoylation levels in different mutants. Tdp1 was depleted by 8h treatment with 1 mM auxin. To reveal PCNA sumoylation, 6His-Pol30 (WT) or 6His-Pol30-K127/164R (2KR) proteins were purified in denaturing conditions on Ni-NTA beads as described in (Davies and Ulrich, 2012). Input or bound fractions were subjected to immunoblotting (IB) with indicated antibodies. (C) The same strains as in (B) were spotted on YEPD media supplemented with indicated amounts of auxin and CPT and imaged 48h post-plating. (D) DNA repair factors identified in the SUMO proteomics analysis. A list of “DNA repair” genes (GO:0006281) was intersected with the two SUMO proteomes described in Figures 3C and 3D. At least 3 unique peptides were identified for each protein shown on the heat map. Values represent relative changes in sumoylation (log2); a cross indicates the absence of unique peptides.

**Supplemental Figure 4.**
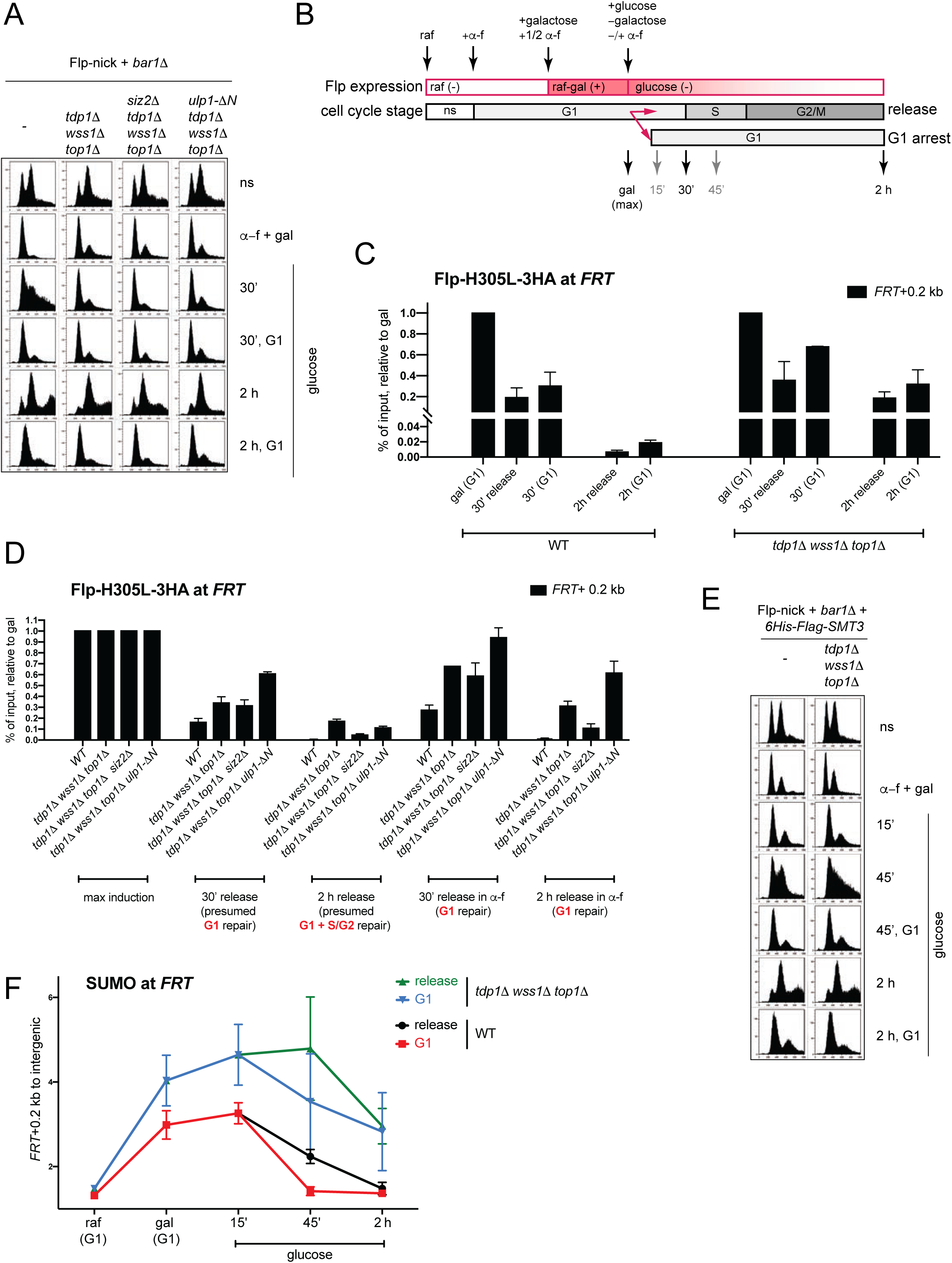
(related to Figure 4) (A) Cell cycle analysis of the cells used in Figures 4E, S4D, and S4E monitored by FACS. (B) Modification of the experimental design used in (Figure 4D) to retain Flp-nick cells in G1. Galactose-induced cultures were split in two before washes. A fraction of yeast cultures was retained in G1 by maintaining *α*-factor. See Methods for details. (C) Comparison of the Flp-nick repair in G1 vs cell cycle release. Yeast cultures were grown as shown in (S4B). ChIP-qPCR without crosslink analysis was performed as in Figure 4E. ChIP-qPCR data are presented as % of inputs adjusted to maximum galactose induction for each genotype. For the release cultures, the data from Figure 4E is used. (D) Comparison of the Flp-nick repair in different mutants in G1 vs cell cycle release. Flp-H305L-3HA levels at *FRT* were monitored as in Figure 4E. The data from Figure 4E and additional G1 retention timepoints were normalized to galactose induction for each genotype. G1 retention was performed as shown in (Figure S4B). (E) Representative cell cycle analyses of cultures shown in Figures 4F and S4F. Cell cycle stage was monitored by FACS. (F) SUMO dynamics at the *FRT* locus. The graph shows the data from Figure 4F and an additional analysis of the *tdp1*Δ *wss1*Δ *top1*Δ mutant.

**Supplemental Figure 5.**
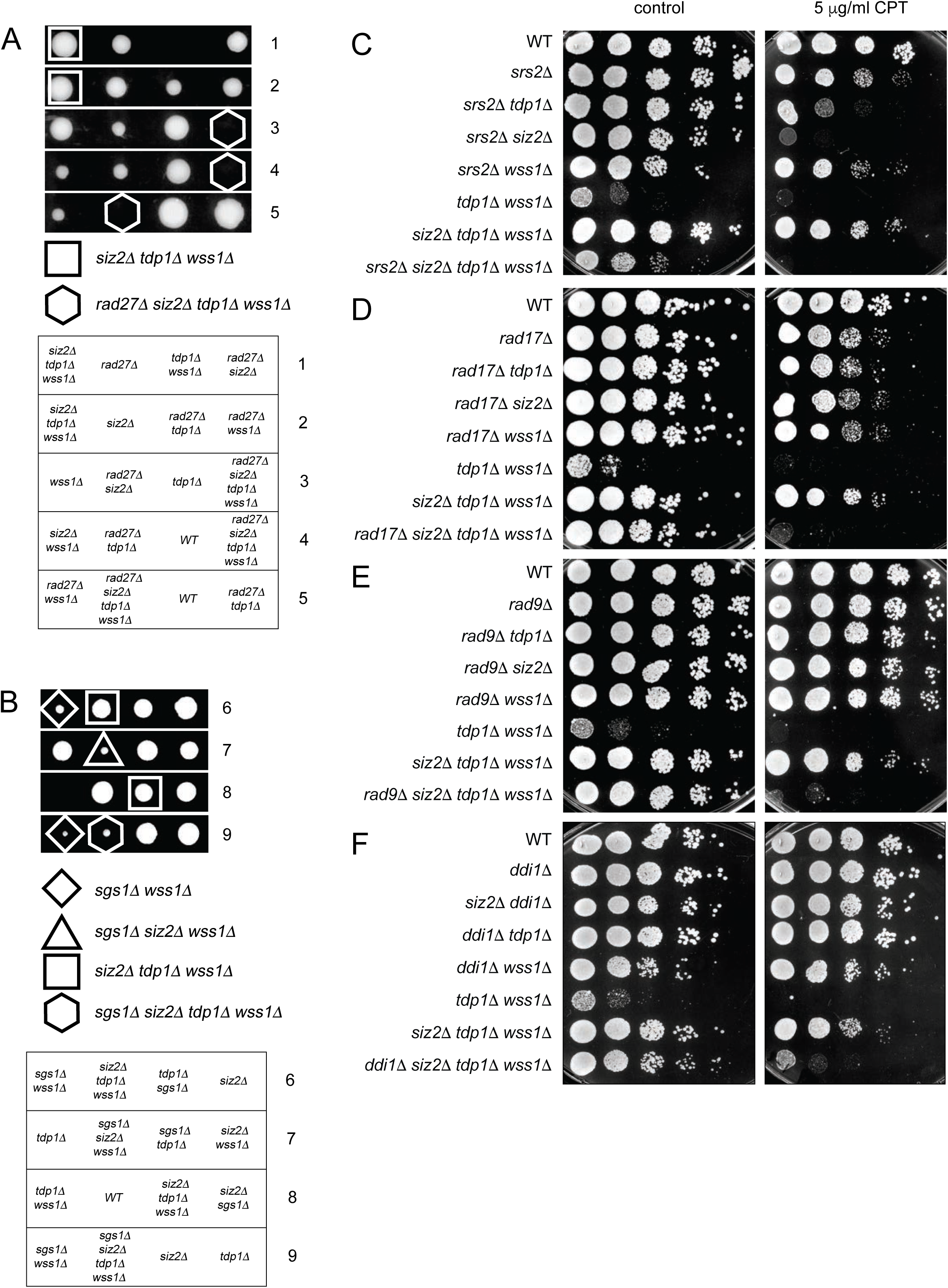
(related to Figure 5) (A) *tdp1*Δ *wss1*Δ *siz2*Δ is synthetic lethal with *rad27*Δ. Tetrad analysis systematically reveals no growth of the combined quadruple mutant. (B) Genetic interactions of *sgs1*Δ combined with *tdp1*Δ, *wss1*Δ, and *siz2*Δ examined by tetrad analysis. (C) - (F) CPT sensitivity of the *tdp1*Δ, *wss1*Δ, and *siz2*Δ mutants combined with *srs2*Δ (C), *rad17*Δ (D), *rad9*Δ (E), or *ddi1*Δ (F). Indicated amount of CPT was added to YEPD medium; images were captured after 48 h of growth.

**Table S1.**
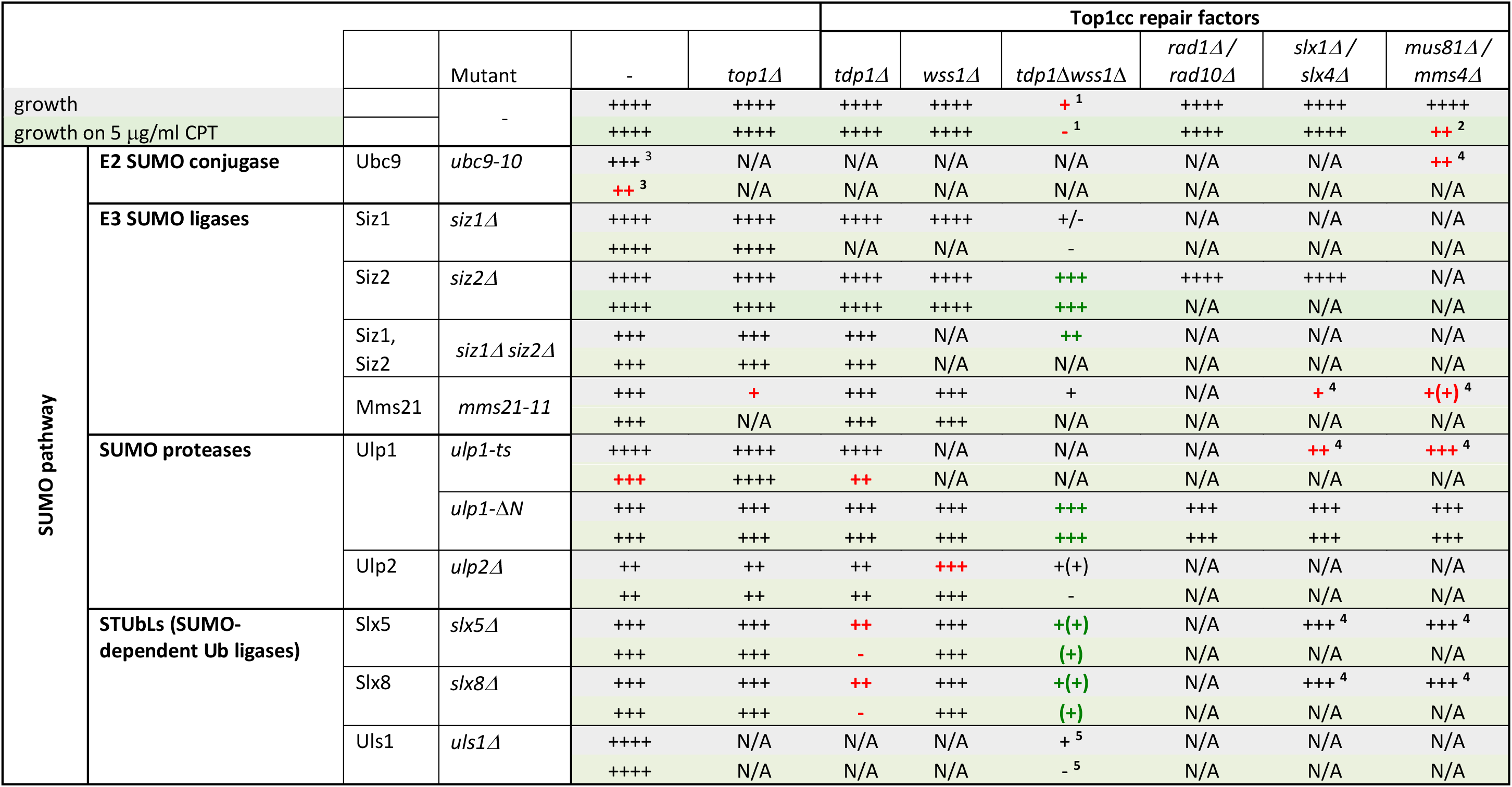
**Genetic interactions and CPT sensitivity of Top1cc repair components and SUMO biogenesis factors.** The table presents relative to wild-type yeast growth rates on complete medium (grey rows) or on 5 μg/ml CPT (green rows) of combined SUMO pathway (left) and Top1cc repair (top) mutants. ++++, WT-like growth; +++, ++, +, negatively affected growth rates; -, no growth; N/A, not tested. Negative genetic interactions are shown in red. Synthetic rescue interactions are shown in green. Genetic interactions documented in other studies: ^1^ (Stingele et al., 2014); ^2^ (Doe et al., 2002); ^3^ (Jacquiau et al., 2005); ^4^ (Costanzo et al., 2016); ^5^ (Sharma et al., 2017).

**Table S2.** **Summary of the extended tetrad analyses of mutants identified in the *tdp1*Δ *wss1*Δ *top1*Δ SATAY genetic screen.** *Growth rates do not reflect CPT sensitivity (quadruple mutants are sensitive to 5 μg/ml CPT). See Figure S5 for spot assays on CPT. ++++, WT-like growth; +++, ++, +, negatively affected growth rates; ±, nearly no growth; –, no growth.

| Process or function |  | WT | *wss1* | *wss1 siz2* | *tdp1 wss1* | *wss1 tdp1 siz2* |
| --- | --- | --- | --- | --- | --- | --- |
|  | WT | ++++ | ++++ | ++++ | ± | +++ |
| HR | *rad52Δ* | +++ | ++ | ++ | - | ++ |
|  | *rad51Δ* | ++++ | ++ | ++ | - | ++ |
| Structural nucleases | *mre11Δ* | +++ | ++ | ++ | - | ++ |
|  | *rad27Δ* | +++ | ++ | ++ | - | - |
|  | *rad10Δ\** | ++++ | ++++ | ++++ | ± | +++ |
|  | *slx4Δ\** | ++++ | ++++ | +++(+) | ± | +++ |
| NHEJ | *yku70Δ* | ++++ | +++ | +++ | ± | +++ |
| DPC protease | *ddi1Δ\** | ++++ | +++ | +++ | - | ++ |
| RecQ helicase | *sgs1Δ* | ++++ | + | + | ± | + |
| Anti-recombinase | *srs2Δ\** | ++++ | ++++ | ++++ | - | ++ |
| DDR checkpoint, 9-1-1 recruitment | *rad17Δ*<br>*rad9Δ* | ++++ | ++++ | ++++ | ± | +++ |

Table S3 – Oligonucleotides, strains, plasmids used in this study and details of their construction.

Data S1 – Results of the SILAC-MS SUMO proteome analysis

Data S2 – Relative transposon coverage in the *tdp1*Δ *wss1*Δ *top1*Δ SATAY genetic screen

## STAR Methods

### RESOURCE AVAILABILITY

#### Lead Contact

Further information and requests for resources and reagents should be directed to and will be fulfilled by the Lead Contact, Françoise Stutz

#### Materials Availability

All unique/stable reagents generated in this study are available from the Lead Contact without restriction.

### EXPERIMENTAL MODEL AND SUBJECT DETAILS

#### Saccharomyces cerevisiae

##### Yeast strains

*Saccharomyces cerevisiae* yeast strains were derived from W303 [*leu2-3,112; trp1-1; can1-100; ura3-1; ade2-1; his3-11,15*] or S288C (BY4741) [*his3*Δ*1; leu2*Δ;*0 lys2*Δ*0; met15*Δ*0; ura3*Δ*0; MATa*] genetic backgrounds. An exhaustive list of strains with respective genotypes is available in Table S3.

##### Yeast growth media

Yeast were grown at 30°C in YEP- or SC- based liquid media or plates supplemented with 20 g/l agar. 2% glucose (default), 2% raffinose, or 2-3% galactose was used as a source of sugar. Selection against the URA3 marker was performed in the presence of 1 mg/ml of 5*-*Fluoroorotic acid (5-FOA). Selection for dominant markers was performed on YEPD-based medium supplemented with 200 μg/ml G418, 200 μg/ml clonNAT or 50 μg/ml Hygromycin B. Prototrophs were selected on SC media lacking the respective amino acids or nucleobases.

YEP medium: 1% yeast extract, 2% peptone.

YEPD medium: YEP, 2% glucose.

SC medium: 1.7 g/l yeast nitrogen base, 5 g/l ammonium sulfate, 2% agar, 2% glucose, 0.87 g/l dropout mix. Dropout mix composition: 0.8 g adenine, 0.8 g uracil, 0.8 g tryptophan, 0.8 g histidine, 0.8 g arginine, 0.8 g methionine, 1.2 g tyrosine, 2.4 g leucine, 1.2 g lysine, 2 g phenylalanine, 8 g threonine.

SD+2% galactose-adenine agar plates used for the transposon screen: 1.7 g/l yeast nitrogen base, 5 g/l ammonium sulfate, 2% agar, 2% galactose, 0.03 g/l isoleucine, 0.15 g/l valine, 0.02 g/l arginine, 0.02 g/l histidine, 0.1 g/l leucine, 0.03 g/l lysine, 0.02 g/l methionine, 0.05 g/l phenylalanine, 0.2 g/l threonine, 0.04 g/l tryptophan, 0.03 g/l tyrosine, 0.02 g/l uracil, 0.1 g/l glutamate, 0.1 g/l aspartate.

#### Escherichia coli

DH5*α E. coli* bacterial strains were grown at 37°C in LB medium or on LB-2% agar plates supplemented with 50 μg/ml of ampicillin for plasmid selection.

### METHOD DETAILS

#### Yeast techniques

##### Construction of yeast strains

*De novo* mutations were introduced into yeast genomes by transformation. Oligonucleotides, template plasmid or genomic DNA used for PCR and transformation are listed in Table S3. Genome editing was verified by colony PCR and/or sequencing. An epitope tag insertion was additionally checked by immunoblotting. Mutants with multiple genomic modifications were obtained by genetic crosses.

##### Yeast transformation

To edit yeast genomic DNA or to transform plasmid DNA (pDNA), log-phase growing yeast cultures were resuspended in LiTE buffer (100 mM LiAc, 10 mM Tris pH 7.5, 1 mM EDTA) and mixed with 100 μg/ml salmon sperm ssDNA, 37.28% [w/v] PEG4000 and purified PCR fragment or pDNA. Cells were incubated for 1-2 h at 30°C, then supplemented with 6% DMSO and a heat shock was performed for 10 min at 42°C. Single colonies were isolated by growth on selective medium.

##### Genetic crosses and tetrad analysis

To perform a yeast genetic cross, two strains of opposite mating type were mixed and grown overnight on rich YEPD medium, selected for diploid markers on corresponding media, and sporulated for 4-5 days on KAC plates (20 g/l potassium acetate, 2.2 g/l yeast extract, 0.5 g/l glucose, 0.87 g/l dropout mix, 20 g/l agar, pH 7.0) until visible tetrads were formed. Asci were digested with 0.5 mg/ml zymolyase 20T, spores were separated by dissection and grown for 2-4 days on rich YEPD media. Genotyping was performed by replica-plating on selective auxotrophic or dominant markers (when available) or made by colony PCR. In some cases, diploids were transformed with pDNA prior to sporulation. The presence of pDNA in colonies formed from spores was tested by growth on selective markers.

##### Colony PCR

Genotyping of yeast and bacterial strains was performed using the Phire Green Hot Start II PCR Master Mix supplemented with the relevant oligonucleotides according to manufacturer’s recommendations.

##### Spot assay

10x serial dilutions of log-phase growing yeast cultures were spotted on agar plates supplemented with 1 mM auxin or indicated amounts of CPT. To induce *pGAL10-Flp-H305L* expression, cells were pre-grown in YEP + 2% raffinose and spotted on 2% glucose- or 2% galactose-containing media.

##### Flp-nick induction for ChIP

Induction of the *pGAL10-flp-H305L* or *pGAL10-flp-H305L-3HA* expression in the Flp-nick or Flp-nick-HA genetic backgrounds (see Table S3 for full genotypes) was performed essentially as described in (Nielsen et al., 2009). Cells were pre-cultured overnight in YEP + 2% raffinose, then diluted to OD_600_=0.15-0.4 in fresh YEP + 2% raffinose media and grown for 3-4 h before galactose induction to enter log phase. Flp-H305L constructs were induced for 2 hours with 3% galactose that was directly added to YEP + 2% raffinose medium. For Flp-nick induction in G1, cells were synchronized for 1-2 h with 200 ng/ml *α*-factor prior to galactose induction; additional 100 ng/ml *α*-factor was added together with galactose. All Flp-nick strains used for G1 arrest contained the *bar1Δ* mutation. To stop galactose induction, two washes using 1/5 yeast culture volume of cold YEP medium were performed. Next, cells were re-suspended in YEP-2% glucose to repress *pGAL1-flp-H305L*. For the glucose release in G1 experiments, 200 ng/ml *α*-factor was added to the YEP medium used for washes, as well as to YEP-2% glucose.

#### Construction of recombinant pDNA and yeast genome editing

Plasmid DNA was constructed either by standard digestion-based cloning or using NEB Builder HiFi DNA Assembly Master Mix. A list oligonucleotides and PCR templates used to generate products for cloning and genome editing is provided in Table S3. Additional details of pDNA construction and genomic DNA modification are available on request.

#### Flow cytometry analysis by FACS

1 ml aliquots of cell cultures at OD_600_=0.3-1 were collected to monitor cell cycle progression by flow cytometry cell sorting (FACS). Cultures were pelleted, re-suspended in 70% EtOH, and stored at 4°C for up to 1-2 weeks. To prepare samples for FACS, fixed cells were centrifuged at 3800 g for 2 min, washed once with 300 μl of NaCi buffer (50 mM NaCi, pH 7.2) and centrifuged for 10 min at 3800 g. Pellets were resuspended in 250 μl of NaCi buffer supplemented with 0.4 mg/ml RNAseA and incubated for 1-3 h at 37°C. DNA was stained with 25 μg/ml propidium iodide for 1 h at 37°C and sonicated 5 times for 5 s with a Bioruptor Twin. Flow cytometric analysis was performed in NaCi buffer using a Gallios flow cytometer. FACS profiles were analyzed by Kaluza software.

#### Chromatin immunoprecipitation (ChIP)

ChIP analysis was performed as detailed in (Serbyn et al., 2020). Flp-nick cultures were grown as described above. 100 ml of yeast cell culture at OD_600_=0.7-1.2 was fixed with 1% formaldehyde for 15 min, neutralized by 250 mM glycine for 5 min at room temperature, kept for 10 min or longer on ice, then pelleted, washed twice with cold 1x PBS (137 mM NaCl, 2.7 mM KCl, 10 mM Na_2_HPO_4_, 2 mM KH_2_PO_4_, pH 7.4) and frozen in liquid nitrogen. Pellets were resuspended in FA lysis buffer (50 mM HEPES-KOH, pH 7.5, 140 mM NaCl, 1 mM EDTA, 1% Triton X-100, 0.1% sodium deoxycholate (w/v)) supplemented with freshly added inhibitor (1 tablet of mini EDTA-free protease inhibitor cocktail / 10ml) and 25 mM N-ethylmaleimide (NEM). Cells were mixed with 500μl glass beads and disrupted in the MagNA Lyser Instrument (5 times for 30 s at 6000 rpm with 1 min pause) at 4°C. The soluble fraction was separated by centrifugation (30 min at 18000 g at 4°C) and further discarded. Chromatin pellets were resuspended in 1 ml of the fresh FA+inhibitor+NEM buffer. DNA was fragmented by sonication in the Bioruptor Twin, 20 cycles of 30 s – 30 s. Lysates were centrifuged for 15 min, 18000 g at 4°C to remove insoluble fraction. Protein concentration in the soluble fraction was defined by the Bio-Rad protein assay. Equivalent quantities of total protein (0.5-1.5 mg) were used for immunoprecipitation with 1 μl of anti-Flag antibody (SUMO ChIPs) or 1 μl of anti-HA (Flp-H305L-3HA ChIPs); 1/10 of each sample (input) was kept separately before mixing with antibody. Immunoprecipitations were performed overnight at 4°C followed by 3 h coupling to 25 μl of washed protein G Dynabeads. IPs were washed once with 500 μl of FA buffer, twice with FA-500 (50 mM HEPES-KOH, pH 7.5, 500 mM NaCl, 1 mM EDTA, 1% Triton X-100, 0.1% sodium deoxycholate (w/v)), twice with Buffer III (10 mM Tris-HCl pH 8, 1 mM EDTA, 250 mM LiCl, 1% IGEPAL, 1% sodium deoxycholate (w/v)) and once with TE (50 mM Tris-HCl pH 7.5, 10 mM EDTA). Complexes were then separated from beads at 65°C by two sequential incubations with 100 μl of elution buffer B (50 mM Tris-HCl pH 7.5, 1% SDS, 10 mM EDTA). The volume of inputs was adjusted to 200 μl with TE buffer. Proteins in ChIP and input samples were digested with 0.75 mg/ml Proteinase K for 2 h at 42°C. Decrosslinking was performed for at least 12 h at 65°C. DNA was purified using either the MinElute PCR purification kit or the Nucleospin Gel and PCR clean up kit (using NTB) according to the manufacturer’s recommendations. DNA was eluted from the column twice with 30 μl sterile H_2_O or the elution buffer from a kit. 2 μl eluates were used for 10 μl qPCR reactions with SYBR Select Master Mix. Oligonucleotide pairs and respective concentrations are listed in Table S3 (FRT+0.2 kb, FRT+0.5 kb FRT+1 kb, intergenic). For all experiments, percent of inputs were calculated. Where indicated, percent of inputs were normalized to the intergenic signal.

#### ChIP without crosslink

Since the *flp-H305L* mutation covalently crosslinks Flp protein to DNA, the formaldehyde crosslinking step was omitted in Flp-H305L-3HA ChIP experiments. Instead, yeast cell cultures were spun, washed once with 1/5 volume of 1xPBS buffer and pellets were directly frozen in liquid nitrogen and stored at –80°C until further processing using the ChIP protocol described above. For Flp-H305L-3HA ChIPs, no NEM was added to the lysis buffer.

#### SATAY library generation

The SATAY transposon screen was performed as in (Michel et al., 2017). Generation of the *tdp1-AID wss1Δ* + auxin library (Figure 1) is described in (Serbyn et al., 2020). The *tdp1*Δ *wss1*Δ *siz2*Δ SATAY library was generated using the strain FSY7292 [*tdp1*Δ*::HPH; wss1*Δ*::HIS3; siz2*Δ*::NATMX6; ade2*Δ*::KANMX6; MATa*] as detailed in (Michel et al., 2017). Briefly, the pBK257-transformed yeast strain was grown in SC-ura+2% raffinose/0.2% glucose to saturation and plated on SD-adenine+2% galactose agar plates. The library was grown for 3 weeks, then yeast colonies were collected, pooled, diluted to an approximate concentration of 2.5×10^6^ cells/ml in 2 liters of SC+2%glucose-adenine medium, grown until saturation, pelleted and frozen. By our estimation, the original library contained approximately 2.2 x10^6^ clones. 500mg of yeast pellet was used to extract genomic DNA, digest with DpnII and NlaIII, circularize, and perform PCR with P5_MiniDs and MiniDs_P7 oligonucleotides. The DNA sequencing library contained equal amounts of DpnII- and NlaIII-digested DNA and was sequenced using the MiSeq v3 chemistry adding 3.4 μl of 100 μM 688_minidsSEQ1210 primer.

The sequencing data were analyzed as described in Michel et al. 2017. The coordinates and number of associated sequencing reads were saved as .bed and .wig files for uploading into a genome browser. The “read_per_gene” value (Figure 1A) takes into consideration the fact that abundant transposons mutants can be detected by multiple sequencing reads. The “tn” value (Figure 5B) counts number of positions per yeast gene body where transposons were inserted and is best suited to identify synthetic sickness genetic interactions (Michel et al., 2017).

#### Protein extraction

To extract proteins, 5-10 ODs of exponentially growing yeast cultures were fixed with 6.25% trichloroacetic acid, kept on ice for 10 min or more, pelleted, washed twice with 100% acetone and dried under vacuum. Dry pellets were resuspended in urea buffer (50 mM Tris-Cl pH 7.5, 5 mM EDTA, 6 M Urea, 1% SDS), mixed with 200 μl of 0.5 mm glass beads and homogenized in the MagNA Lyser 5 times for 45 s at 4°C. Lysates were then incubated for 10 min at 65°C and centrifuged for 10 min at 18000 g to remove foam. Lysates were mixed with 1.5x volume of sample buffer (3% SDS, 15% glycerol, 0.1 M Tris pH 6.8, 0.0133% bromophenol blue, 0.95 M 2-mercaptoethanol) and boiled for 10 min.

#### Immunoblotting

Protein extracts were resolved by SDS-PAGE using 6-14% gels, transferred to nitrocellulose or PVDF membranes, blocked with 5% dry milk dissolved in TBS-T (150 mM NaCl, 20 mM Tris-HCl, 0.05% Tween, pH 7.4). Primary and secondary antibodies were diluted in TBS-T containing 5% dry milk as indicated in the Key Resource Table. Immunoblot signals were revealed with WesternBright ECL or Sirius HRP substrate; images were taken with the Li-COR Odyssey Imaging System and quantified using Fiji.

#### SUMO assay

To isolate SUMO proteins, the genomic copy of yeast *SMT3* was N-terminally tagged with 6His-Flag. Exponentially growing yeast cultures were fixed with 5% trichloroacetic acid, cooled on ice for approximately 1 h, pelleted, washed twice with 100% acetone and dried under vacuum. Yeast pellets were re-suspended in 1 ml of freshly prepared G-buffer (100 mM sodium phosphate pH=8, 10 mM Tris-HCl pH=8, 6M guanidinium, 10 mM 2-mercaptoethanol, 0.1% Triton X-100, 5 mM MG132, 25 mM NEM, EDTA-free protease inhibitor cocktail), mixed with 500 μl of 0.5 mm glass beads and homogenized in the MagNA Lyser 5 times for 45 s at 4°C. Recovered lysates were centrifuged for 20 min, 18000 g at room temperature. The concentration of proteins in the soluble fraction was measured by the Bio-Rad protein assay. Total proteins (3-5 mg in 1 ml of buffer) were mixed with 80 μl of Ni-NTA agarose pre-washed twice with water and once with G-buffer and rotated on a wheel for 2 h at room temperature. After binding, Ni-NTA resins were washed once with 500 μl of G-buffer, three times with fresh U-buffer (8 M urea, 100 mM sodium phosphate pH=6.4, 10 mM Tris-HCl pH=6.4, 10 mM 2-mercaptoethanol, 0.1% Triton X-100, EDTA-free protease inhibitor cocktail). Bound proteins were next eluted from agarose beads by 5 min boiling with 40 μl 3x sample buffer (125 mM Tris-HCl pH=6.8, 4% SDS, 286 mM 2-mercaptoethanol, 0.02 mg/ml bromophenol blue, 20% glycerol). To prepare input fractions, 40 μl of total cell lysates were diluted 10 times in water, supplemented with 10% trichloroacetic acid, incubated 30 min, centrifuged at room temperature for 30 min, 18000 g and re-suspended in sample buffer. Ni-NTA bound and input fractions were further analyzed by immunoblotting.

The sumoylation status of 6His-PCNA was evaluated as described previously (Davies and Ulrich, 2012).

#### Purification of sumoylated proteins and SILAC-MS analysis

The enrichment of SUMOylated proteins in preparation for MS analysis was performed as previously described (Albuquerque et al., 2013) but with the following exceptions: Each mutant strain was grown in 1-liter of synthetic media containing either light or heavy stable isotope-labeled lysine and arginine until an OD_600_ of 0.3, at which point auxin (1 mM final concentration) was added to deplete Tdp1. Cells were harvested following 6 hours of auxin treatment and lysed under denaturing condition as previously described (Albuquerque et al., 2013). Following lysis, protein concentrations were determined by Bradford Reagent (Bio-Rad) and were normalized prior to mixing and subsequent purification of SUMOylated proteins. Nano-flow LC-MS/MS analysis was performed on a Thermo Scientific Ultimate 3000 Nano-LC System and a Thermo Scientific Orbitrap Fusion Lumos mass spectrometer; acquired via NIH S10 OD023498. Data analysis for SILAC-labeled samples was performed as previously described, with the exception that all proteins were required to have a minimum of 3 unique peptides (Suhandynata et al., 2019). A list of “DNA repair” genes (GO:0006281) was extracted from the R package “org.Sc.sgd.db”, v.3.7.0.

## KEY RESOURCES TABLE

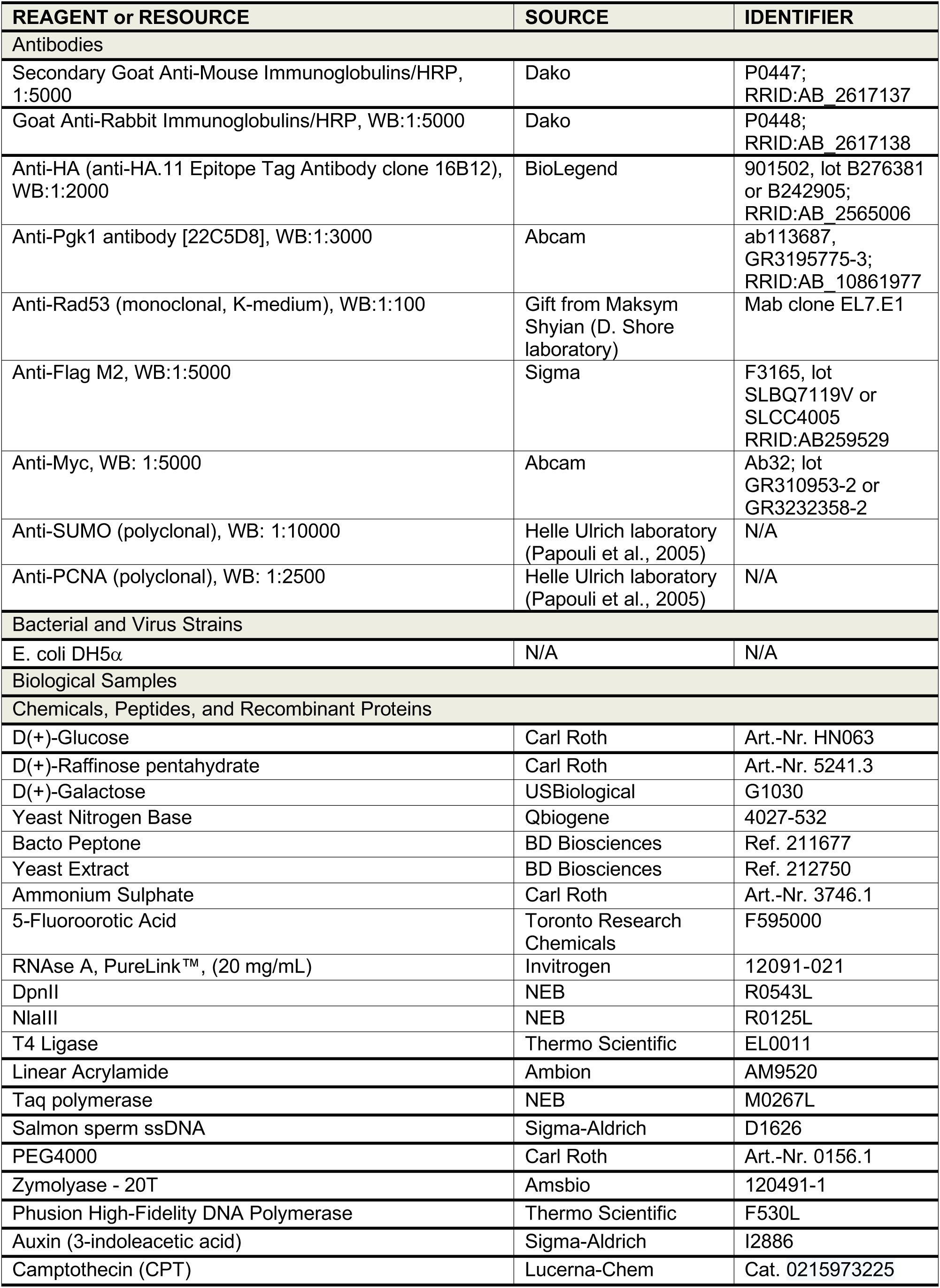
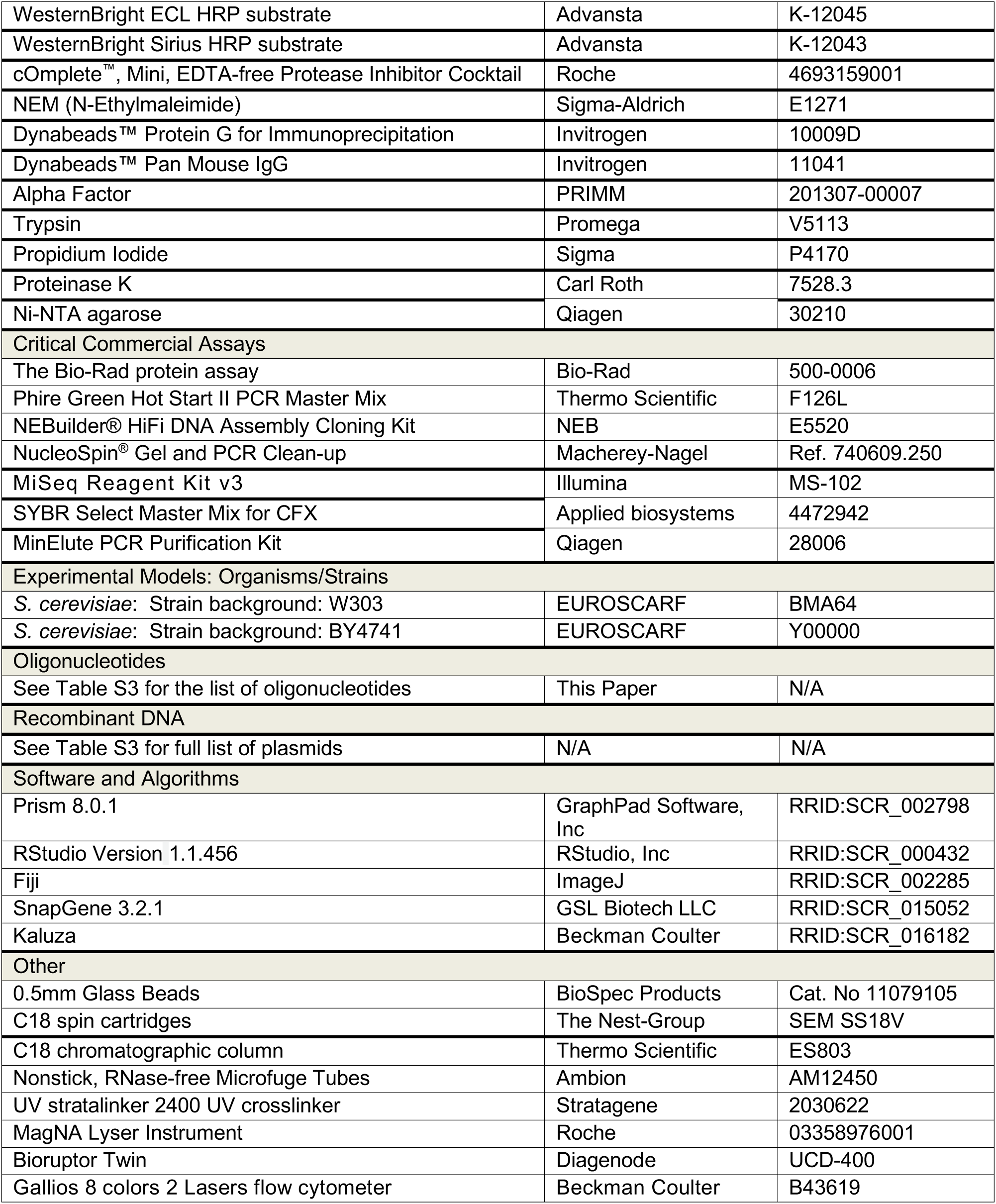

